# Identification of novel human derived influenza viruses in pigs with zoonotic potential

**DOI:** 10.1101/2021.06.08.447649

**Authors:** Rodrigo Tapia, Bárbara Brito, Marco Saavedra, Juan Mena, Tamara García-Salum, Raveen Rathnasinghe, Gonzalo Barriga, Karla Tapia, Victoria García, Sergio Bucarey, Yunho Jang, David Wentworth, Montserrat Torremorell, Víctor Neira, Rafael A. Medina

## Abstract

In 2009, a novel swine influenza A virus (IAV) emerged causing a global pandemic that highlighted the role of swine as a reservoir. To date, there is limited information about swine IAV circulating in Latin America. We identified two swine H1N2 and one divergent swine H3N2 viruses that co-circulated in Chilean swine together with the 2009 H1N1 pandemic strain (A(H1N1)pdm09). Phylogenetic analysis revealed several human-to-swine IAV introductions occurring as early as the mid-1980s, and since 2009, several introductions of the A(H1N1)pdm09 strain. Antigenic cartography confirmed that these viruses were antigenically unique and identified drifted variants within the clusters. Human sera from the Chilean general population showed an age dependent mid to low-level antibody mediated protection against swine H1N2 and A(H1N1)pdm09-like viruses and a poor protection against the swine H3N2 virus, highlighting the zoonotic potential of this strain. Our results underscore the epidemiological importance of studying swine IAV in Latin America for epidemic and pandemic preparedness.

## Introduction

Influenza A virus (IAV) circulates endemically in nature in hosts such as birds, dogs, horses, swine, and humans, among others, representing a constant concern to both public and animal health worldwide. Swine have an important role in the ecology of IAV since they are susceptible to both avian and human strains and have been proposed as a “mixing vessel” for the generation of novel reassortant strains and an intermediate host for interspecies transmission. While the emergence of the 2009 H1N1 influenza A pandemic (A(H1N1)pdm09), first detected in Mexico, occurred due to the reassortment in swine of viruses previously circulating in swine, birds and humans, it must be noted that no specific evidence has shown that swine contributed to the generation of older pandemics (Novel Swine-Origin Influenza et al., 2009; Smith et al., 2009). Nevertheless, early on during the A(H1N1)pdm09 outbreak in humans, the disease was also subsequently detected in commercial pigs in several countries (Corzo et al., 2013; Nelson et al., 2012; Pereda et al., 2010; Watson et al., 2015), highlighting the role of swine as an important reservoir for the introduction of human strains that might further reassort and generate viruses with zoonotic potential. The scarce sequence data available at that time evidenced the significant gap of knowledge of the IAVs circulating in swine and underscored the need for conducting systematic surveillance studies in Latin America.

IAV is one of the most frequently detected pathogens in commercial swine farms, resulting in important economic losses. The most widely distributed subtypes present globally in swine are H1N1, H1N2 and H3N2. These viruses show a high level of genetic and antigenic diversity within each subtype, and several lineages have been reported in different geographical regions worldwide (Anderson et al., 2016; Lorusso et al., 2011; Nelson et al., 2015b; Nelson et al., 2019; Vincent et al., 2014; Watson *et al*., 2015).

The genetic diversity of IAV is due to the high mutation rate inherent to the replication of this virus, and its ability to reassort different segments of its genome. This occurs when two or more IAVs infect cells of the same host, sometimes combining virus genes adapted to different host species. This can generate antigenically novel IAVs by a process called ‘antigenic shift’, due to the reassortment of segments encoding the surface hemagglutinin (HA) and neuraminidase (NA) glycoproteins. In addition, ‘antigenic drift’ can occur due to the appearance of mutations in the glycoproteins because of immune selection pressure. Changes in the HA can cause major changes in the antigenicity of the virus(Medina and Garcia-Sastre, 2011).

In Chile, intensive swine farms represent >95% of pig production, and pork is the second most produced meat in the country, with an industry that has been growing steadily in the last decade (Neira et al., 2017a). Nevertheless, limited information is available on the IAVs circulating in swine, and only a small number of HA genes have been identified as human-like IAV strains (Nelson et al., 2015a). To determine the origins and evolutionary history of these swine IAVs and to elucidate their genomic and antigenic diversity, a thorough sampling and in-depth analyses of the circulating viruses are needed. In addition, to better understand the inter-species transmission dynamics and to determine the zoonotic potential of these swine IAV strains, it is important to conduct risk assessment studies in pertinent populations to evaluate the presence of cross-reactive antibodies in humans against these swine viruses.

Here we performed a systematic swine and human molecular epidemiological study to determine the diversity and origin of swine IAVs circulating in commercial swine farms in Chile. We identified three novel swine viral genotypes, which were generated by at least three reassortment events, resulting in an H3N2, and two antigenically distinct H1N2 viruses. These viruses are related to human viruses from the late 1980s to mid-1990s but contain the internal genes of the A(H1N1)pdm09 pandemic strain. Chilean swine IAVs are genetically distinct from previously reported IAV. Antigenically these viruses are different from other swine and human strains. The general human population showed a mid to low-level antibody protection against the endemic H1N2 strains, but a significantly lower protection against the H3N2 virus, which highlights this as a potential zoonotic strain.

## Results

### Swine IAVs circulate endemically in commercial swine farms in Chile

To determine whether IAV was circulating in different commercial farms and to what extent, we conducted a cross-sectional serological study in three different production systems in Chile (Fig. 1a). We determined that some animals had detectable antibodies against IAV during the first ∼30 days of life, which were possibly maternally derived antibodies transferred during lactation. However, by 14-15 weeks of age (∼100 days) all three farms showed a marked increase in antibody prevalence in the animals, which was indicative of active IAV infections and confirmed that the virus was endemic in these swine populations (Fig. 1b-d). Given that the peak antibody response occurs approximately 28 days after infection, this also suggested that swine infections began to occur 3-4 weeks earlier, at 11-12 weeks (∼80 days) of age once maternal antibodies had begun to wane. Hence, to perform active surveillance we obtained samples from 8-11 week (∼56-70 days) old animals with and without clinical signs suggestive of influenza infection.

**Figure 1.**
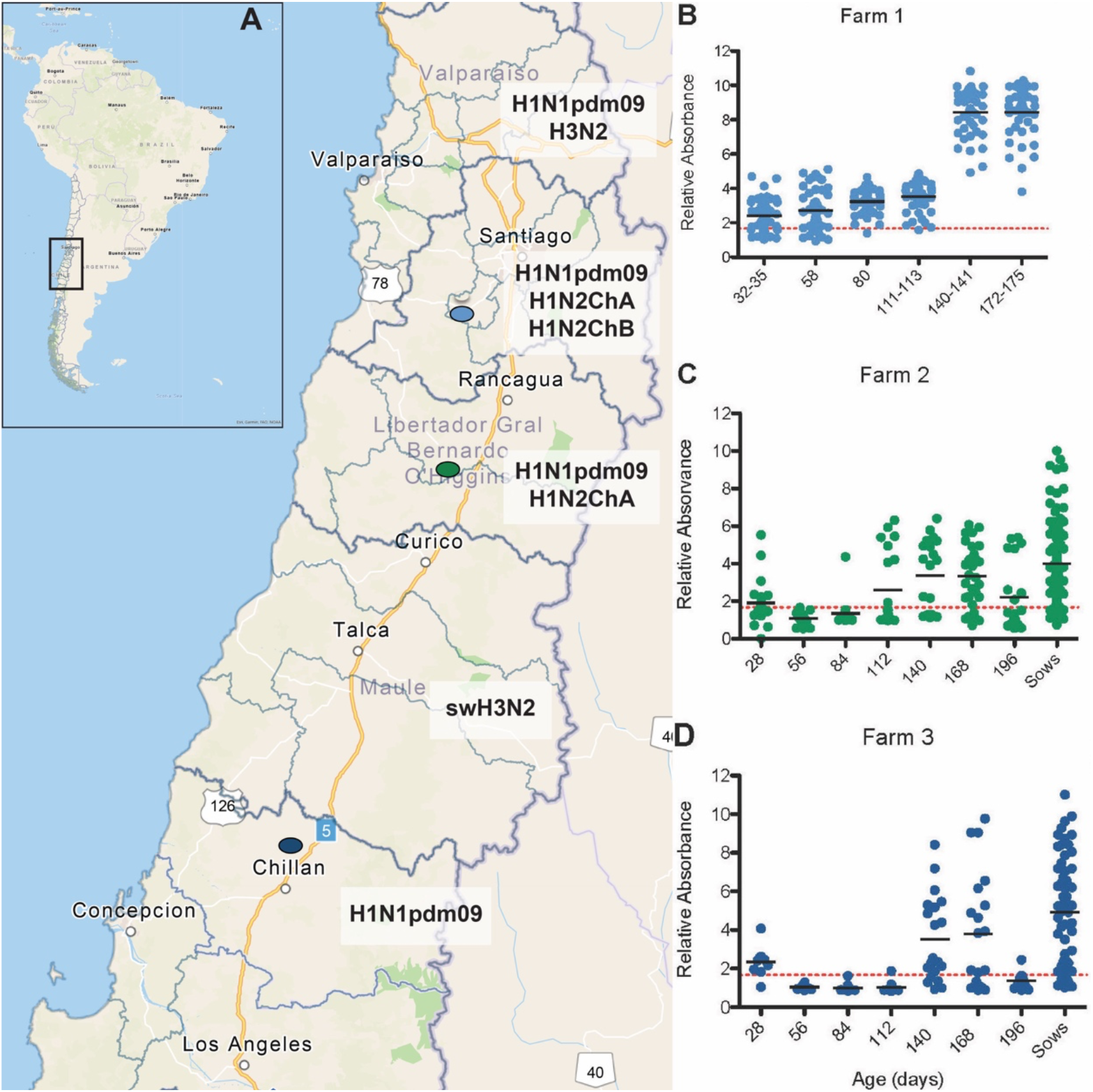
Surveillance sites and serological evidence of endemic circulation of IAVs in Chile. **(A)**. Map of South America and magnification of central regions of Chile indicating the geographical distribution of IAV subtypes identified by administrate region. The north to south depiction of viral strains found are shown per administrative region: Valparaiso region, A(H1N1)pdm09 and H3N2; Metropolitan region A(H1N1)pdm09, H1N2ChA and H1N2ChB; O’Higgins region A(H1N1)pdm09 and H1N2ChA; Maule region, H3N2 and Ñuble region, H1N1p. **(B-D)** Serological profiles of IAV on three commercial swine farms located in 3 different administrative regions of Chile as indicated in the map. Maps were generated using ArcGIS Online tool, sources: Esri, Here, Garmin, Fao, NOAA and USGS.

To obtain representative samples of the swine IAV circulating in commercial swine farms in Chile, we sampled 39 farms representing 93.8% of the industrialized farms in the country. We collected a total of 1,237 samples, including 1,046 (84.6%) nasal swine Abs, 188 (15.2%) oral fluids and 3 lungs (0.2%). Of these, 235 (22.5%) nasal swine Abs, 68 (36.2%) oral fluids, and 3 lungs (100%) tested positive to IAV, which resulted in a total of 306 (24.5%) positive samples. Twenty-eight out of 39 (72%) farms tested positive to IAV at least once during the study. Positive IAV farms were found in all geographical regions sampled (Fig 1a).

### High diversity and multiple reassortant IAVs circulate in swine in Chile

To gain a comprehensive understanding of the viral diversity in the country, we sequenced a total of 70 IAV isolates (52 whole genome sequences and 18 HA sequences) from 26 farms. Of the 70 HA sequences analyzed, 68 belonged to the H1 subtype and two were of the H3 subtype. We identified eight distinct swine IAV genotypes (Fig. 2). Phylogenetic analyses revealed the circulation of two predominant subtypes, two novel and distinct lineages of the swine H1N2 subtype, and the A(H1N1)pdm09-like subtype. The HA sequences of the swine H1N2 viruses were grouped into two major groups that were highly divergent, belonging to two monophyletic clusters with up to 14% nucleotide differences between them (Fig. 3a). These HA segments are of human-origin, but highly distinct from the δ cluster swine IAV previously described in North America. For these new clusters, we propose to name them Chilean H1 A (ChH1 A) and Chilean H1 B (ChH1 B) lineages. All other H1 HAs (27/68) were classified into an A(H1N1)pdm09-like cluster (Fig. 3a). In addition, we isolated two previously uncharacterized swine H3N2 viruses (Fig. 2) circulating in two different farms, which were divergent from all other known swine and human viruses (Fig. 3b)

**Figure 2.**
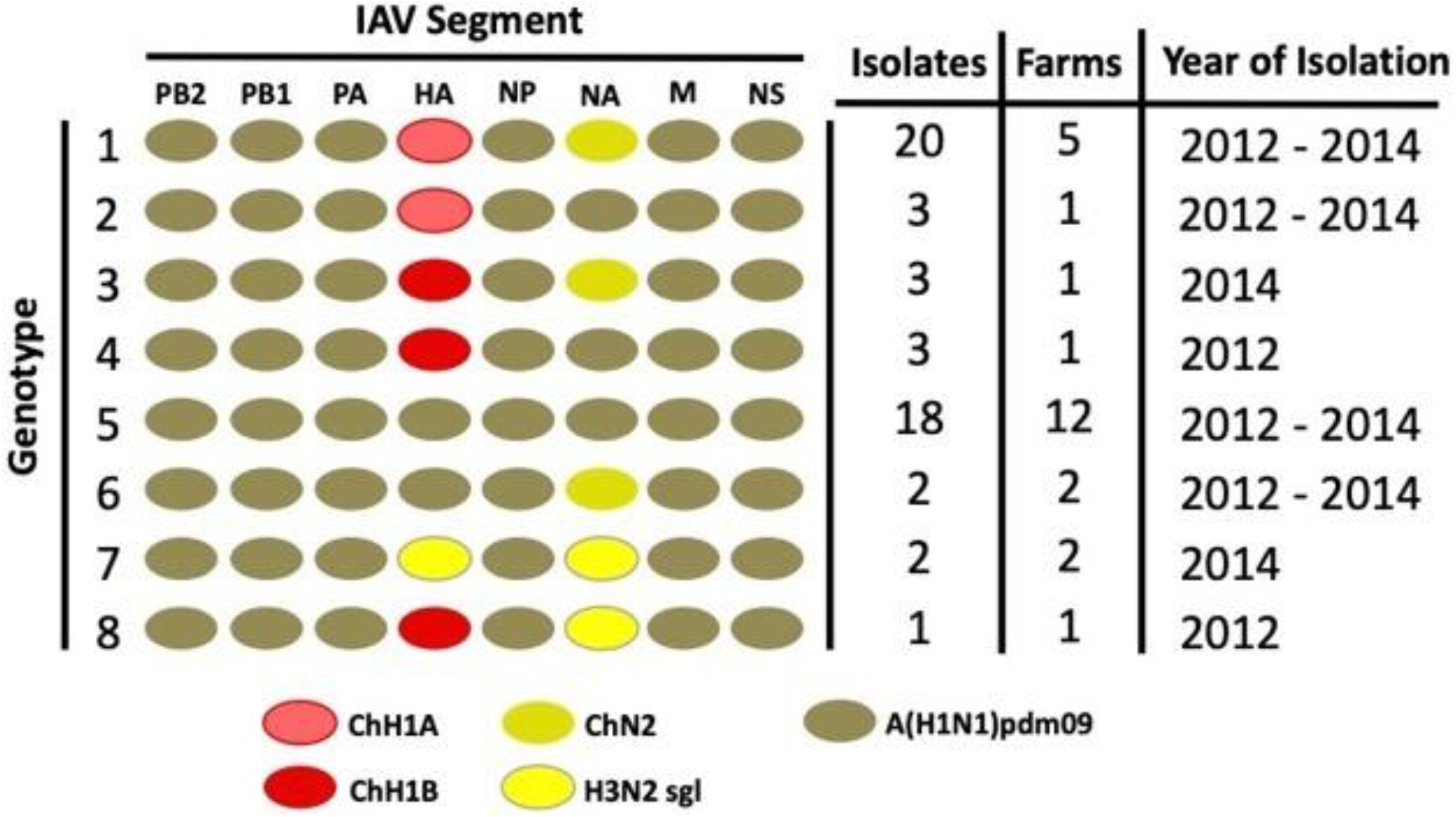
Genotype analyses of novel swine IAV found in Chile. Genetic makeup and lineages of virus subtypes found in swine from commercial farms in Chile during 2012 to 2014. The left column lists the genotype number, and each oval shape represents one of the eight IAV segments corresponding to the indicated lineages. The right-hand side columns indicate the number of times the genotype was isolated, from how many farms and the year of identification of each genotype. ChH1A, Chilean H1A cluster; ChH1B, Chilean H1B cluster; ChN2, Chilean N2 cluster, H3N2 singleton virus; and A(H1N1)pdm09, pandemic H1N1 2009 –like viruses. Segment abbreviations indicate: PB2, Polymerase Basic protein 2; PB1, Polymerase Basic protein 1; PA, Polymerase Acidic protein; HA, Hemagglutinin; NP, Nucleoprotein; NA, Neuraminidase; M, Matrix protein and NS, Non-Structural protein.

**Figure 3.**
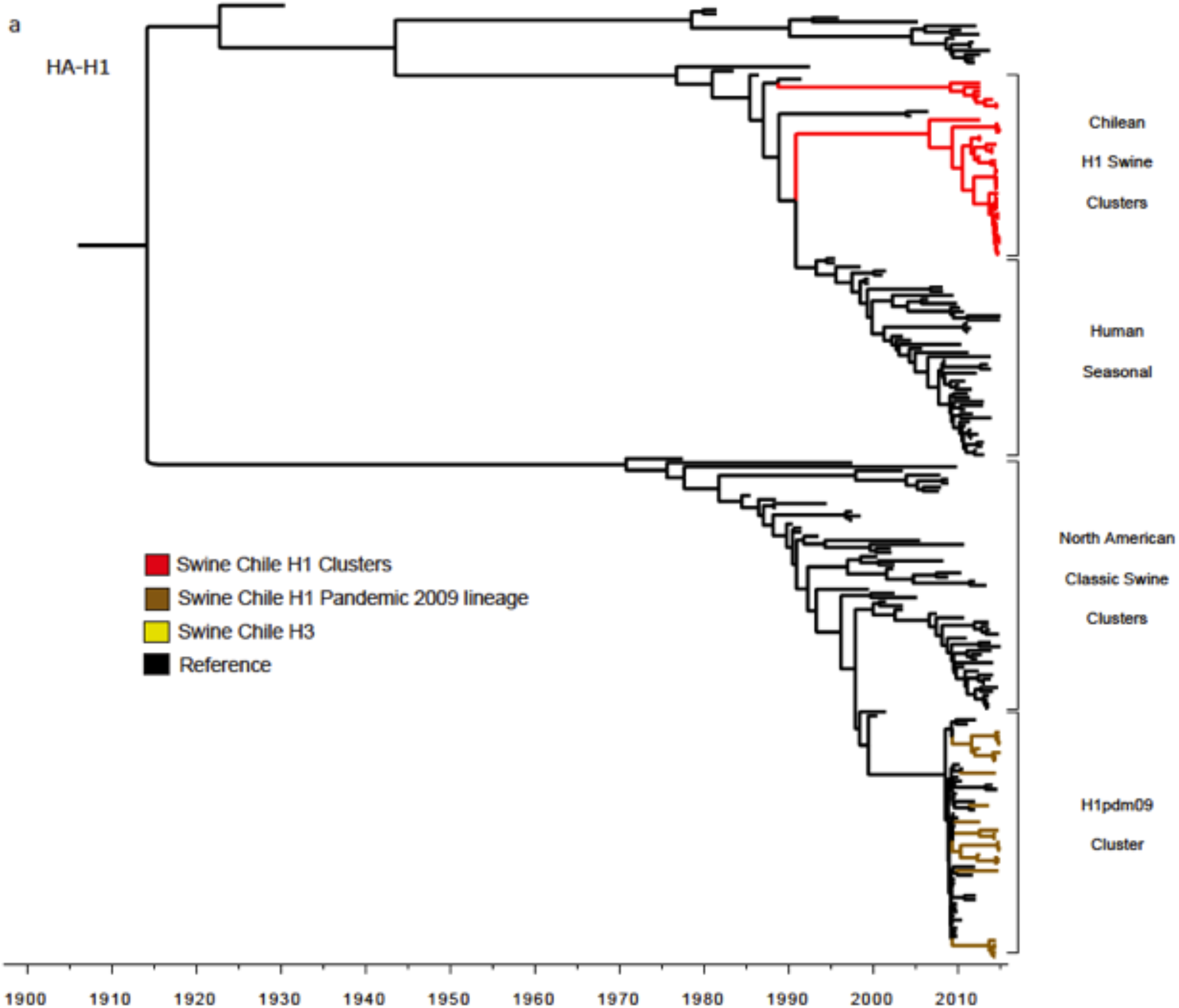
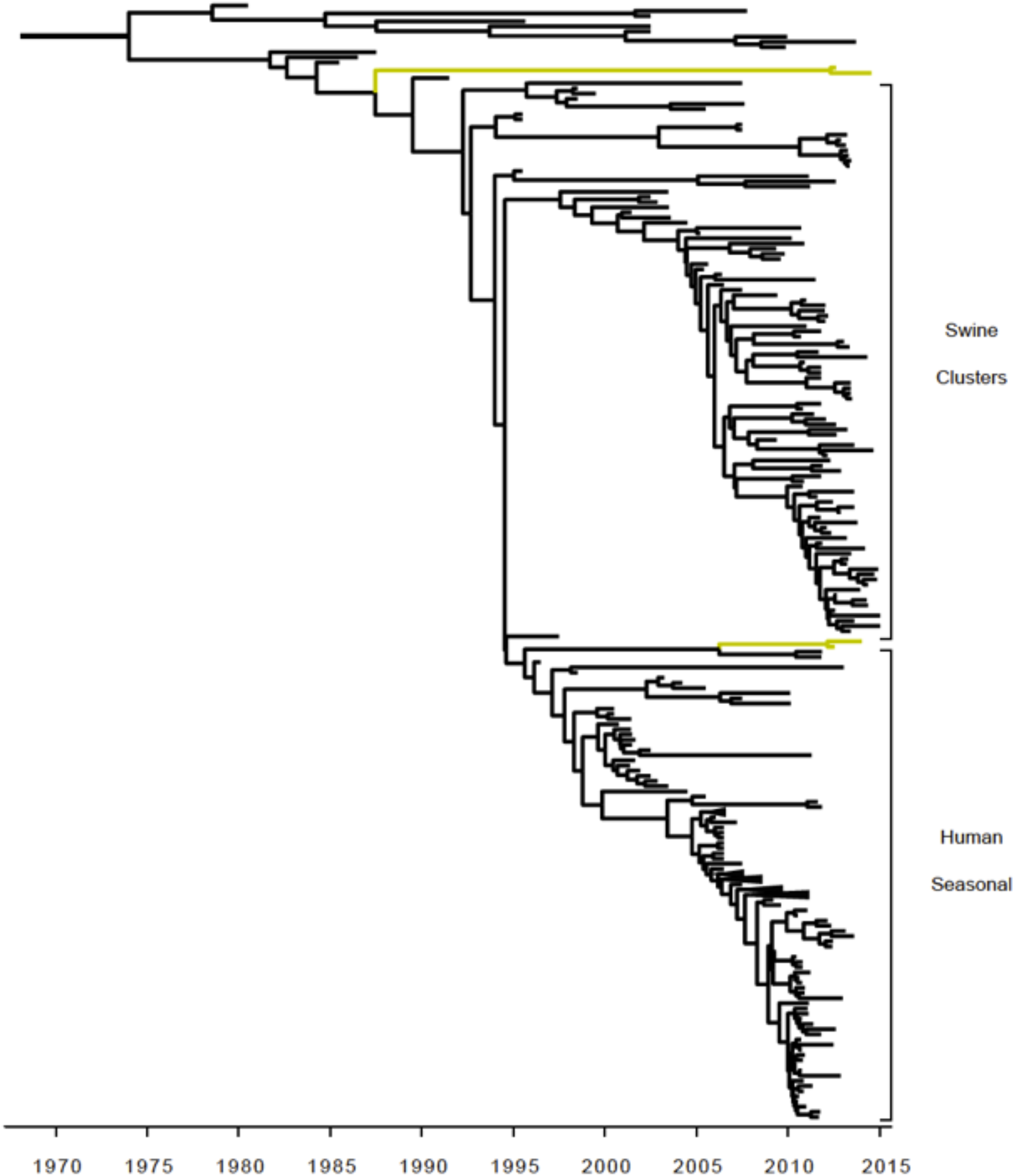
Novel H1 and H3 swine influenza viruses identified in Chile. Maximum clade credibility trees depicting TMRCA estimates, reconstructed using HA gene segment of influenza viruses collected from human and swine. **(A)** Tree reconstructed using H1 type influenza viral reference sequences from human and swine viruses collected globally. Branches of the tree containing the viruses from the newly described endemic Chilean swine viruses are highlighted in red, whereas viruses from the swine A(H1N1)pdm09 (collected in Chile) are highlighted in brown. **(B)** Tree reconstructed using H3 viruses collected from human and swine globally. Viruses collected from Chilean swine are highlighted in yellow.

The HA of most of the swine H1N2 viruses belonged to cluster ChH1A (34/41). The most closely related virus was a human IAV from 2000 (A/Chile/4795/2000(H1N1)). Analyses of the time to the most recent common ancestor (TMRCA) of all the viruses from this cluster that is shared with a human virus from Chile, was estimated to be May of 1994 (95% highest posterior density interval [HPD], July 1990 – August 1998). In addition, the TMRCA representing the viruses that belong to the ChH1A cluster was estimated at August 2006 (95% HPD November 2000 – September 2009) (Fig. 4a). The viruses from ChH1 B cluster were more distant to the δ clusters, suggesting that these viruses could be related to human viruses that originated from the δ swine cluster in North America. The branching of cluster ChH1B from ChH1 A was estimated to be November 1988 (95% HPD January 1986 – December 1990), and the TMRCA of all the viruses in the ChH1 B cluster was estimated at February 2009 (95% HPD March 2004 – November 2010) (Fig. 4a). Overall, the phylogeny suggests that both clusters are of human origin, which are derived from viruses that were introduced into pigs 20-32 years ago and that they became endemic within the Chilean swine population during those years. Based on the newly proposed designation of swine IAV both clades are classified as human seasonal lineage 1B.2 (Anderson *et al*., 2016).

**Figure 4.**
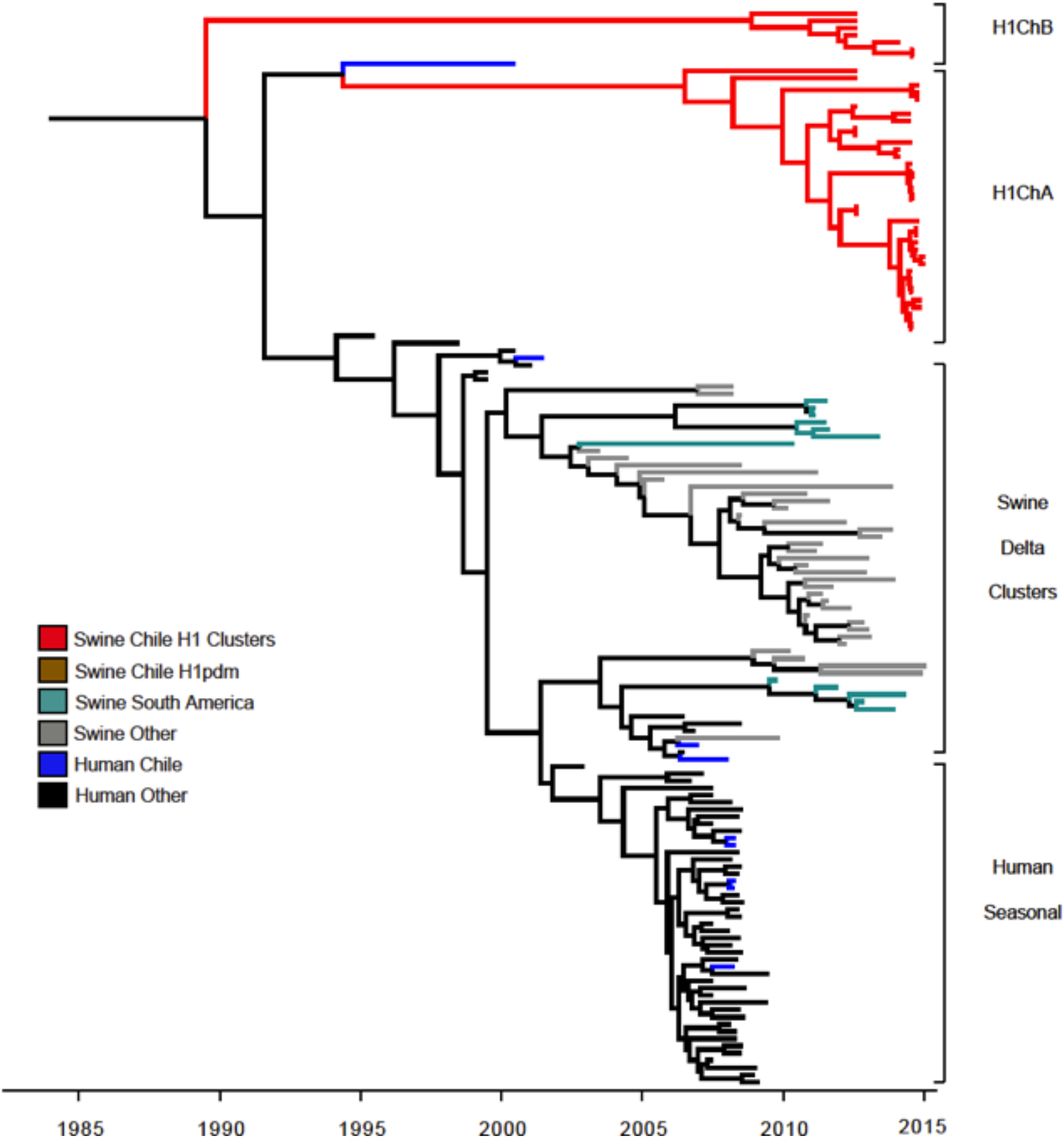
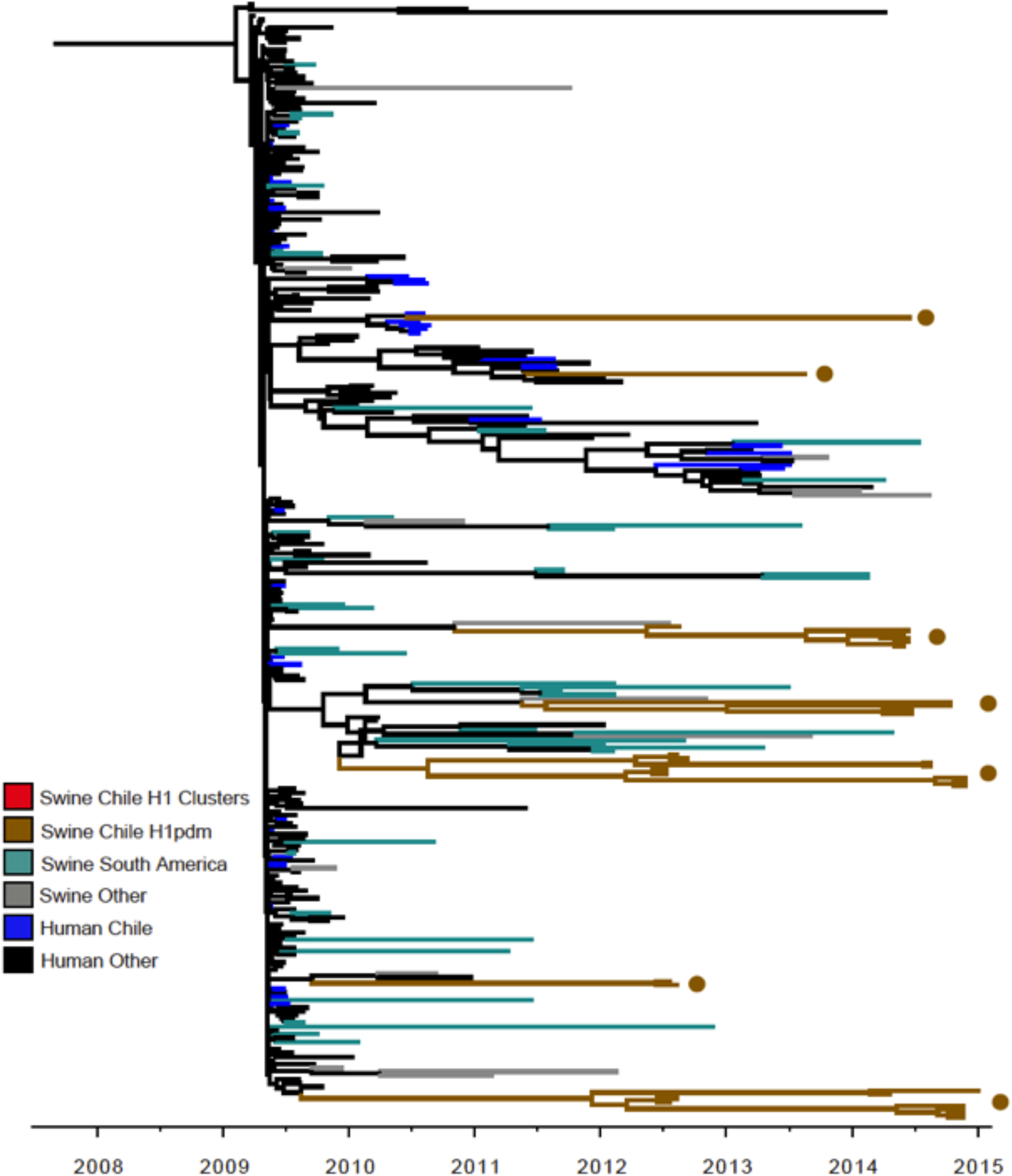
Two divergent H1N2 swine influenza viruses derived from old human seasonal viruses co-circulate with swA(H1N1)pdm09-like strains. Maximum clade credibility tree constructed depicting TMRCA estimates of HA gene of H1 type influenza viruses. **(A)** H1 viruses and reference viruses only from the human seasonal/delta swine/Chilean endemic swine (branches highlighted in red) clusters (excluding pandemic and classic swine viruses). Two clusters of viruses circulating endemically in Chile (ChH1Ch A and ChH1 B) are observed, with a related Chilean human influenza virus. **(B)** H1 maximum clade credibility tree constructed using only viruses belonging to the A(H1N1)pdm09 IAV lineage. Viruses collected from Chilean swine are highlighted in brown. Sampled viruses are interleaved with other reference human viruses, including sequences obtained in Chilean human, suggesting at least seven different introductions from human to commercial swine (*). For figures A and B viruses collected from human in Chile and other parts of the world are highlighted in blue and black, respectively. Viruses collected from swine in South America are in blue-green color, and viruses collected from swine in other parts of the world are highlighted in grey.

### Introductions of A(H1N1)pdm09 from human to swine

To elucidate the diversity and time of introduction of the swine A(H1N1)pdm09-like viruses, we isolated and obtained the full genome sequences of 114 A(H1N1)pdm09 viruses from infected humans during 2009 to 2013 in Chile. The phylogeny of the A(H1N1)pdm09-like viruses showed an interleaved pattern of viruses collected from both species, suggesting at least seven independent introductions of these viruses into swine farms after the 2009 H1N1 human pandemic (Fig. 4b). Additionally, there were two A(H1N1)pdm09 HA introductions that were closely related (genetically) to viruses obtained from infected individuals in Chile from the previous year, suggesting recent reverse zoonotic events. This demonstrates a dynamic human-swine transmission of the A(H1N1)pdm09-like IAVs, which are highly disseminated in the swine population, as they were found in 12 different farms that were not geographically related.

The H3 viruses were related to 2 human-like H3 IAVs singletons reported recently, which are not related to the I-IV North American swine clusters previously described (Fig. 3b)(Nelson et al., 2015d). The TMRCA of one H3 was estimated to be April 2006 (95% HPD September 1998 - January 2009). The second H3 virus was highly divergent compared to any published sequence elsewhere; and had an estimated TMRCA as early as June 1987 (95% HPD May 1990 – June 1984) (Fig. 3b). Each H3 was found in a single farm, which suggests that they were introduced and maintained throughout this time and may be only present in a few sites.

In the case of the NA genes, they belonged to either the N1 (24/52) or N2 (28/52) subtypes. All N1 segments sequenced were derived from the human A(H1N1)pdm09-like N1 cluster (Fig. 5a). Their phylogeny shows the N1 sequences grouping together with other human and swine origin A(H1N1)pdm09 sequences, also suggesting at least seven recent independent introductions into the swine population, like the A(H1N1)pdm09 HA sequences (Fig. 5b). From them, at least three A(H1N1)pdm09 NA introductions are directly related to Chilean human isolates. As with the HA, the A(H1N1)pdm09-like N1 was found in 12 geographically distant farms, suggesting that these genotypes are maintained independently in different production systems. Most N2 segments (89%, 25 out of 28) belonged to a monophyletic cluster suggesting a single introduction of this segment that became widespread via reassortment and evolved within the swine population (Fig. 5c). The branching of this cluster from other swine N2 genes occurred in May 1992 (95% HPD December 1988 – July 1994), while the most recent common ancestor shared among these sequences was September 2007 (95% HPD January 2002-November 2010). These N2 segments are very distinct compared to others circulating in swine elsewhere, hence we propose to name it Chilean N2 (ChN2). There were three additional N2 sequences corresponding to two (singletons), which were not part of the major cluster, and were not related among them (two of them were associated with the H3 sequences, Fig. 5c). TMRCA of these singletons and their closest reference sequences was February 1989 (95% HPD July 1983-September 1992) and February 2001 (95% HPD March 1997-December 2005), respectively.

**Figure 5.**
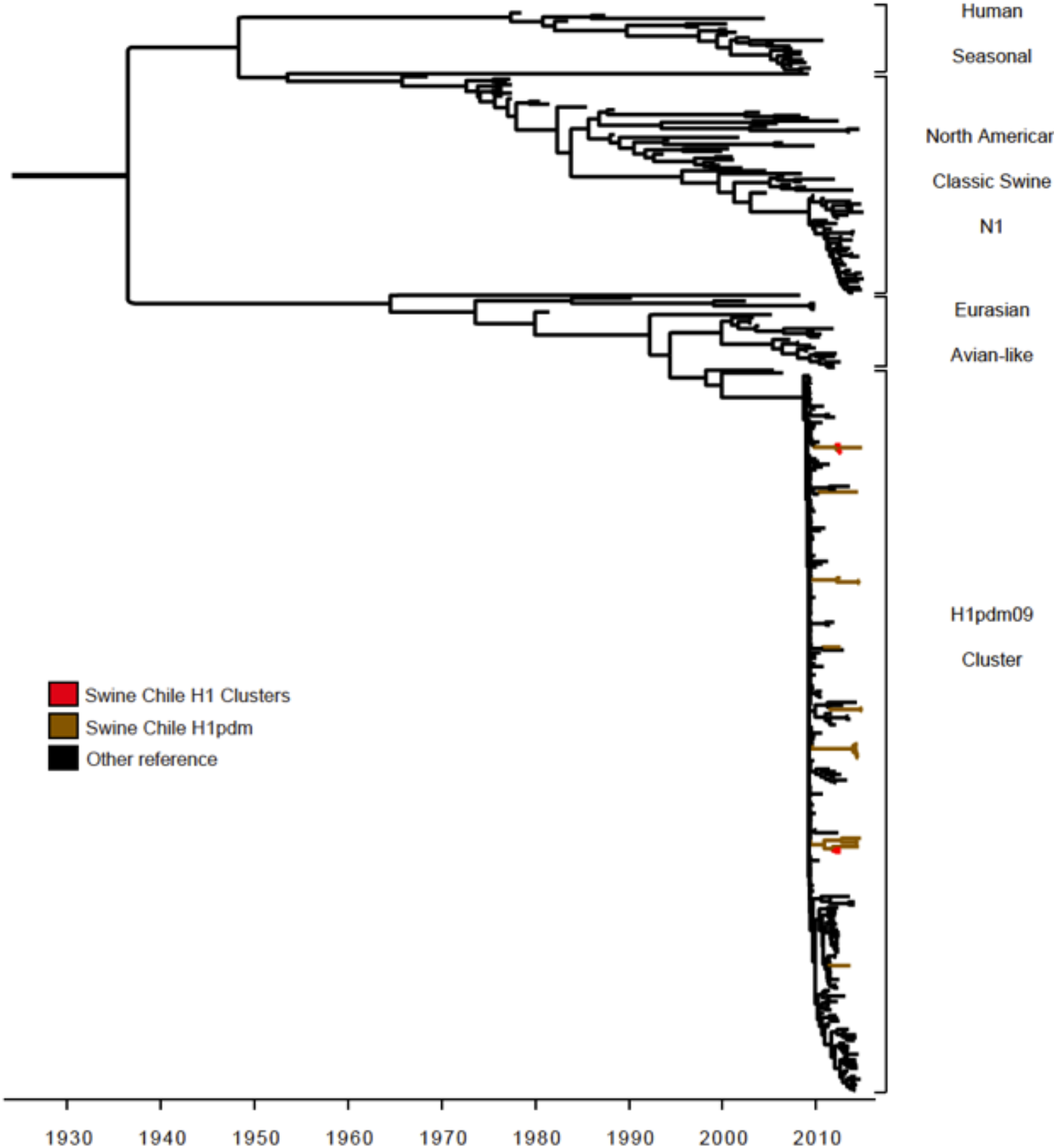
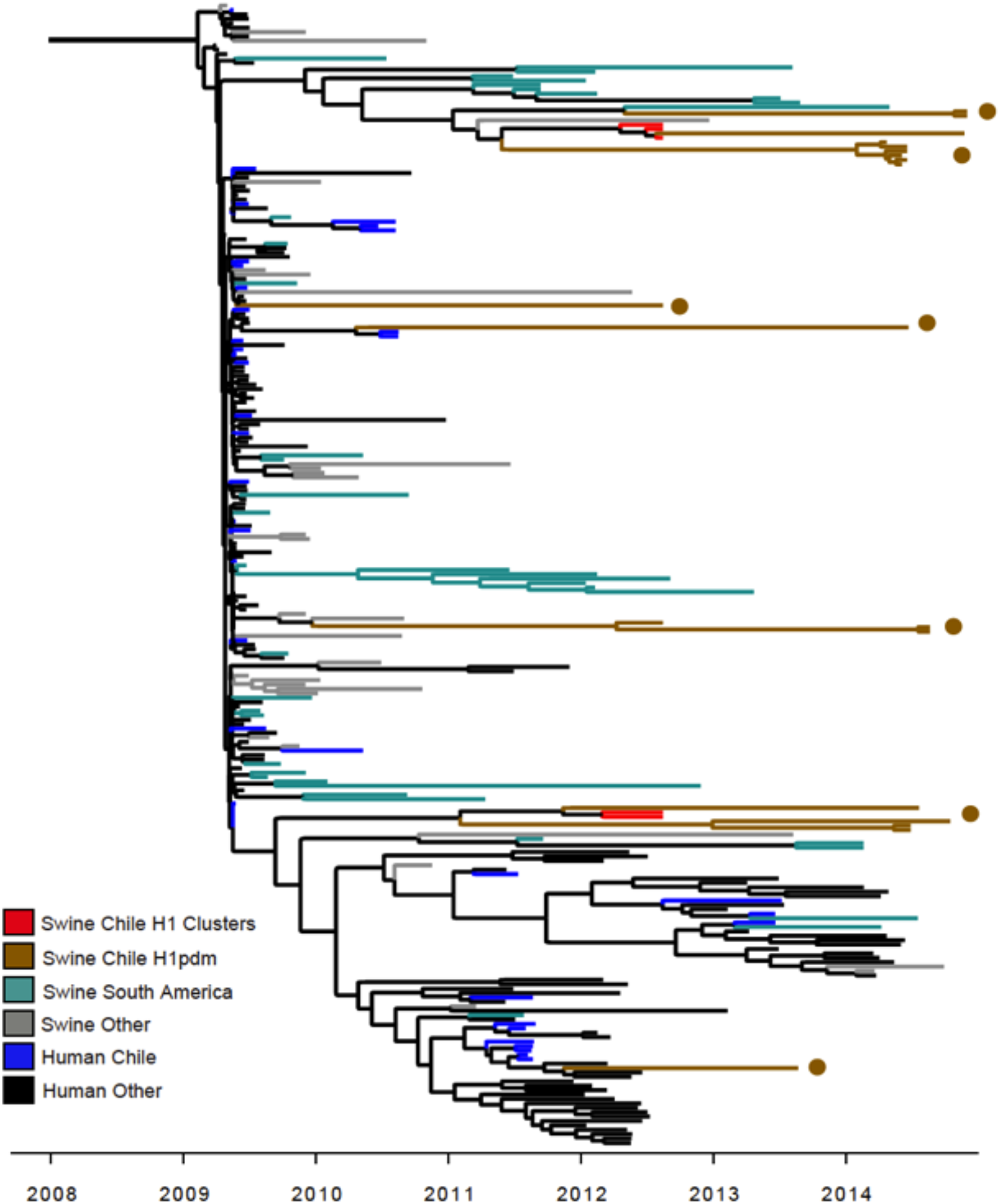
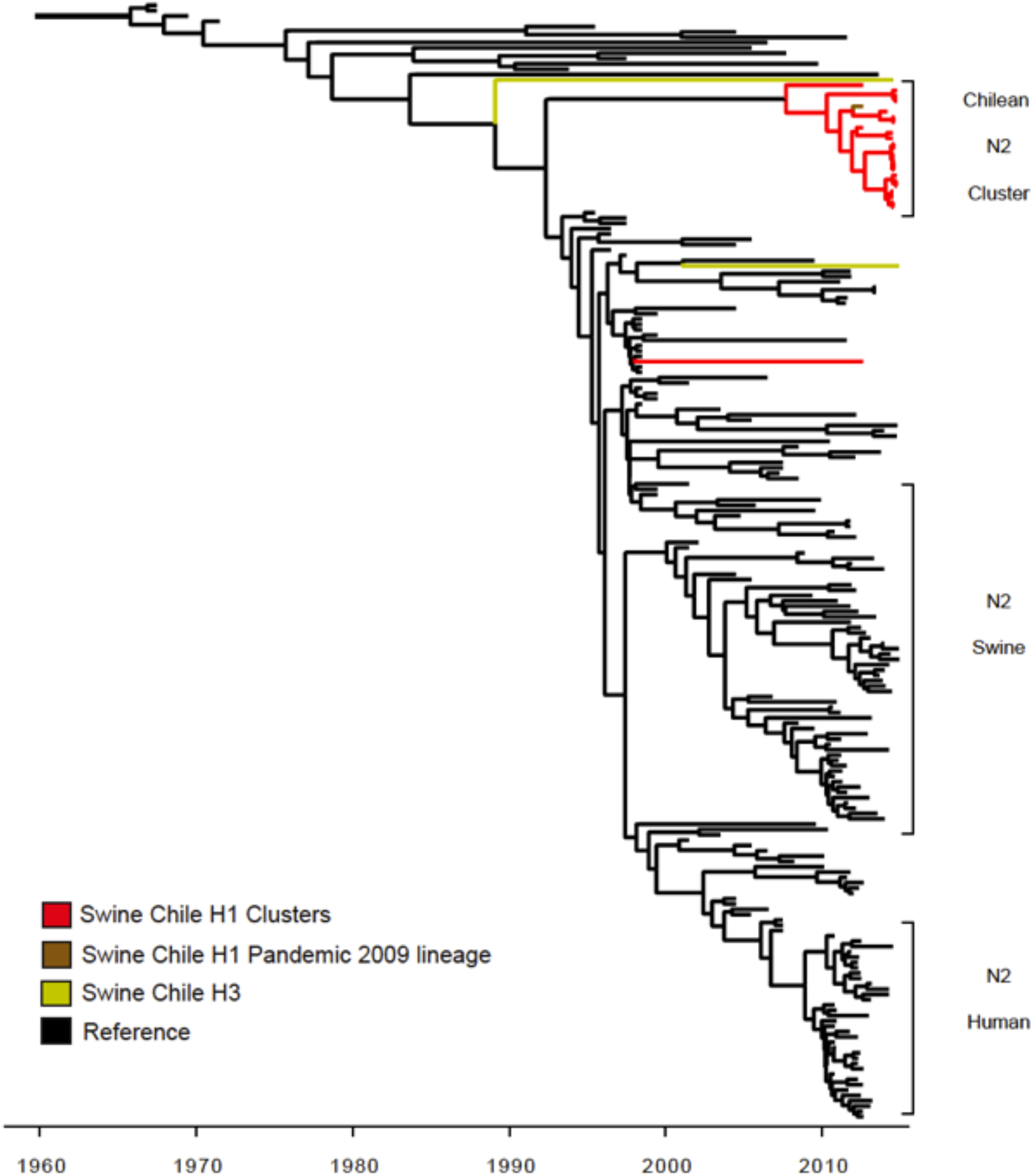
The N1 segments belong to the swA(H1N1)pdm09-like lineage and N2 genes group in a major distinct cluster. Maximum clade credibility trees depicting TMRCA estimates of NA genes segment of influenza viruses collected from human and swine. **(A)** N1, four related N1 lineages are shown: human, swine classic, Eurasian avian-like and N1 pdm. Virus are colored according to the HA original clustering (described for Fig. 3 and 4). **(B)** A higher resolution of the swA(H1N1)pdm09-like lineage is depicted. **(C)** N2, viruses are colored according to the HA original clustering (described for Fig, 3 and 4). Most of the viruses having the ChH1 endemic strain belong to a ‘N2 Chilean cluster’. Viruses having a H3 type HA were divergent; one of them is a singleton sharing an early common ancestor with N2 Chilean viruses. The remaining virus N2 (that correspond to a H3 type) was related to ancestors of the N2 swine and human viruses.

### Human derived A(H1N1)pdm09 internal genes are present in all Chilean swine IAVs

Analysis of the internal genes revealed that all the IAV isolates contained the internal segments derived from the A(H1N1)pdm09-like strain (Fig. 6 and 7, Supplementary Fig. 1). Interestingly, of the 52 matrix (M) segment sequences, 49 were found in a monophyletic cluster (TMRCA May 2010, 95% HPD April 2009 - March 2011), which is related with early human A(H1N1)pdm09 isolates from Chile in 2009 (Fig. 6). Similarly, the phylogeny of PA and NP genes showed that most of the viruses are grouped into one to three major clusters from 2009 - 2010 (Supplementary Fig. 1a and b). This suggests that M, PA and NP segments were introduced into the swine population early after the 2009 pandemic outbreak in Chile and have remained mainly fixed in the swine population since then. In contrast, the phylogeny of the NS, PB1 and PB2 genes showed multiple introductions of these pandemic genes (Fig. 7, and Supplementary Fig. 1c and d). Overall, these results suggest that independent introductions occurred, possibly simultaneously, in different farms in the country generating multiple reassortment events that gave rise to the different genotypes observed.

**Figure 6.**
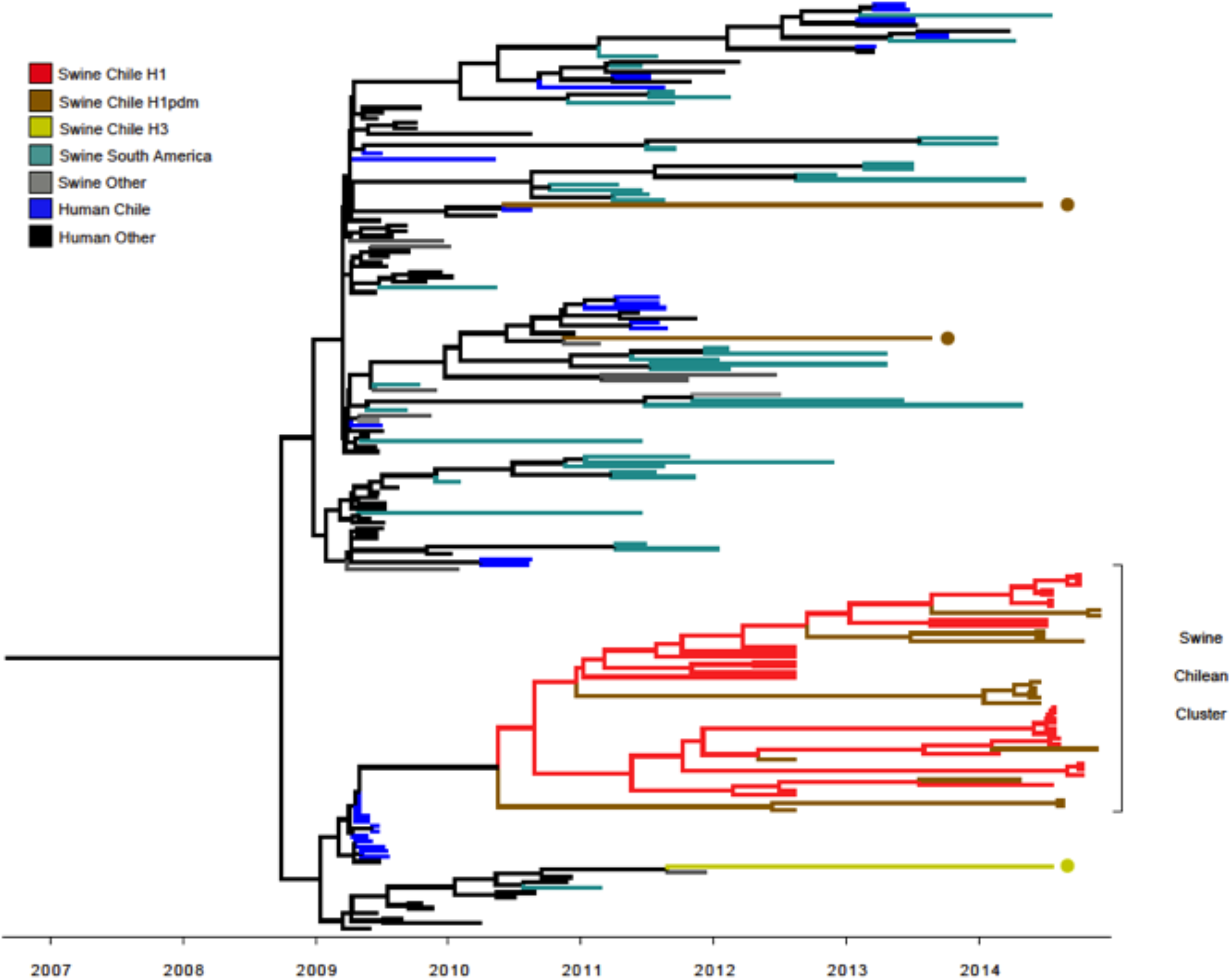
The M Segment genes derives from early A(H1N1)pdm09 human strains. Maximum clade credibility tree depicting TMRCA estimates of M gene segment of influenza viruses collected from human and swine. Most viruses ChH1 and A(H1N1)pdm0-like were grouped within the same cluster and had a most recent common ancestor with viruses from the human A(H1N1)pdm09. The M segment of three swA(H1N1)pdm09-like viruses and from the H3 influenza virus were grouped with other influenza viruses.

**Figure 7.**
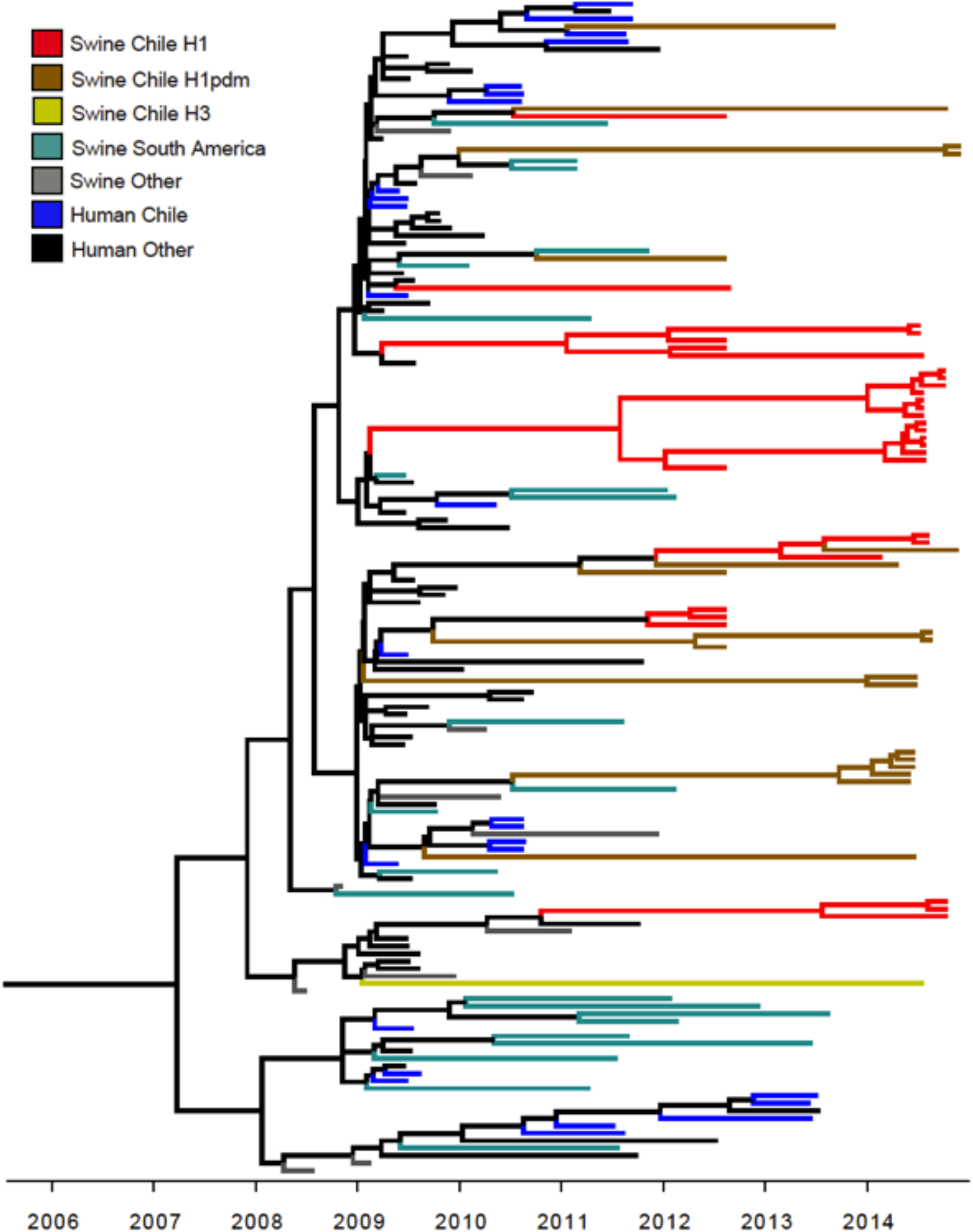
The NS A(H1N1)pdm09 gene of the swIAVs has been introduced multiple times in the swine population. Maximum clade credibility tree depicting TMRCA estimates of NS gene segment of influenza viruses collected from human and swine. Viruses from the ChH1 and A(H1N1)pdm09-like and H3 are interleaved with reference viruses from human and swine from other countries. The NS segments associated to Chilean H1 (red), H1pdm (green), and Chilean H3 (yellow) HA segments are shown. There have been multiple introductions of the NS pandemic gene into the swine population since the human 2009 H1N1 pandemic.

### The novel swine H1ChA, H1ChB and A(H1N1)pdm09-like genetic clusters are antigenically distinct

Due to the divergent characteristic of the ChH1A, ChH1B found in Chile and the multiple introductions of the A(H1N1)pdm09 HA, we analyzed their antigenic relatedness by antigenic cartography. There was no cross-reactivity between the ChH1A and ChH1B viruses, and as expected these viruses were also vastly different from the swine A(H1N1)pdm09-like viruses. This indicates that the novel Chilean genetic clusters also correspond to antigenically distinct clusters (Fig. 9). In general, we observed a high level of cross-reactivity among strains from the same genetic cluster; however, we identified six ChH1 A and three H1pdm09 viruses that were antigenically divergent from their respective antigenic clusters (Fig. 8), suggesting the emergence of potential drift viruses in some farms.

**Figure 8.**
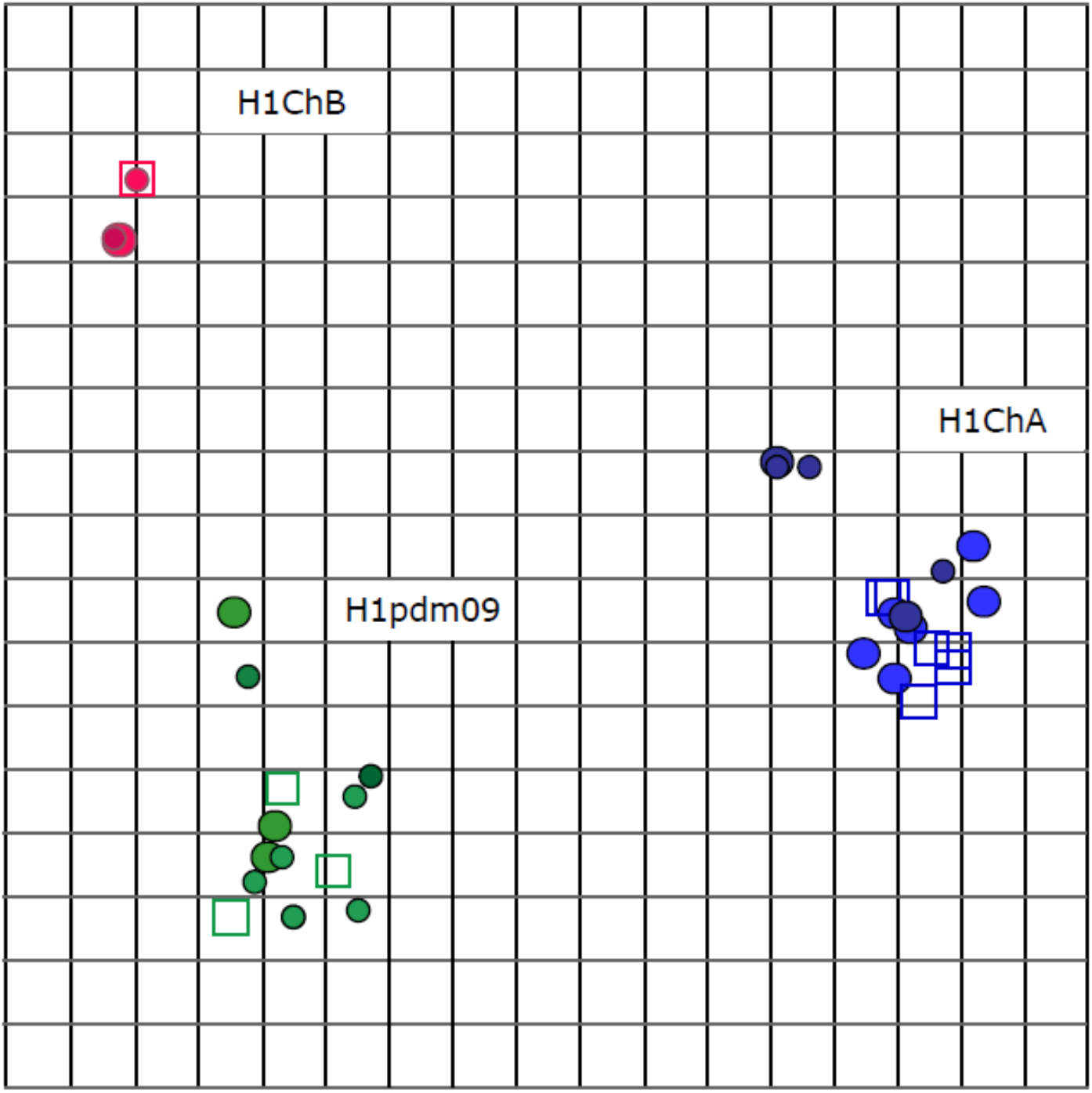
The novel ChH1N2A and ChH1N2B viruses are antigenically distinct. Antigenic map constructed with HIA titers from ChH1A, ChH1B and A(H1N1)pdm09-like clusters. Circles correspond to antigens and squares correspond to antisera. Color represents the antigenic cluster to which the strain belongs; H1ChA is blue, H1ChB is red, and A(H1N1)pdm09 is green. Clusters were identified by a k-means clustering algorithm. The vertical and horizontal axes both represent antigenic distance, and the orientation of the map within these axes is free. The spacing between grid lines is 1 unit of antigenic distance, corresponding to a 2-fold dilution of antiserum in the HIA.

### The swine IAVs are phylogenetically distinct from contemporary human viruses and the general population has limited cross-protective antibodies against them

To determine the relationship of the newly identified swine IAVs to contemporary human seasonal viruses and to the World Health Organization recommended human vaccine strains of the last 40 years, we performed phylogenetic analyses with an extensive list of human viruses isolated since 1918 to 2015 and the respective recommended vaccine strains. Analyses performed with the ChH1A, ChH1B, swine H3 and the ChN2 genes confirmed their unique identity and confirmed that these strains are substantially different from previous identified viruses (Supplementary Fig. 2a-c). Importantly, these swine viruses clustered distantly to all previously recommended vaccines groups

Due to the novel genomic and antigenic properties of the ChH1A and ChH1B viruses, the unusual origin of the swine H3N2 strains, and to determine if the swine A(H1N1)pdm09-like antigenic properties are different from human strains, we performed a risk assessment and evaluated if humans have cross-reactive antibodies against these novel swine viruses. Proper evaluation of cross-protective immunity involves the assessment of the local populations that could become exposed to these viruses. Hence, we used 237 human sera collected, through an influenza clinical cohort in Chile in 2009 to 2015. These individuals were born from 1915 to 2015, covering the entire time frame of potential exposure to the contemporary human influenza viruses of the last 100 years (Fig. 11a). We observed a moderate to high level of protection (titers of neutralizing antibodies) against the ChH1N2A, ChH1N2B and swine A(H1N1)pdm09-like viruses in middle-aged individuals (date of birth: 1965-94; Fig. 11b). In contrast, lower reactivity was observed in the youngest and oldest individuals, born in 1995-2015 and 1915-1944, respectively (individuals < 20 and > 70 years of age, Fig. 11). This pattern of ‘age-related protection’ was not observed for the swine H3N2 viruses, where similar low to mid-levels of antibody reactivity were seen for individuals of all ages (Fig. 10b), highlighting this virus as a higher zoonotic potential risk to humans

**Figure 10.**
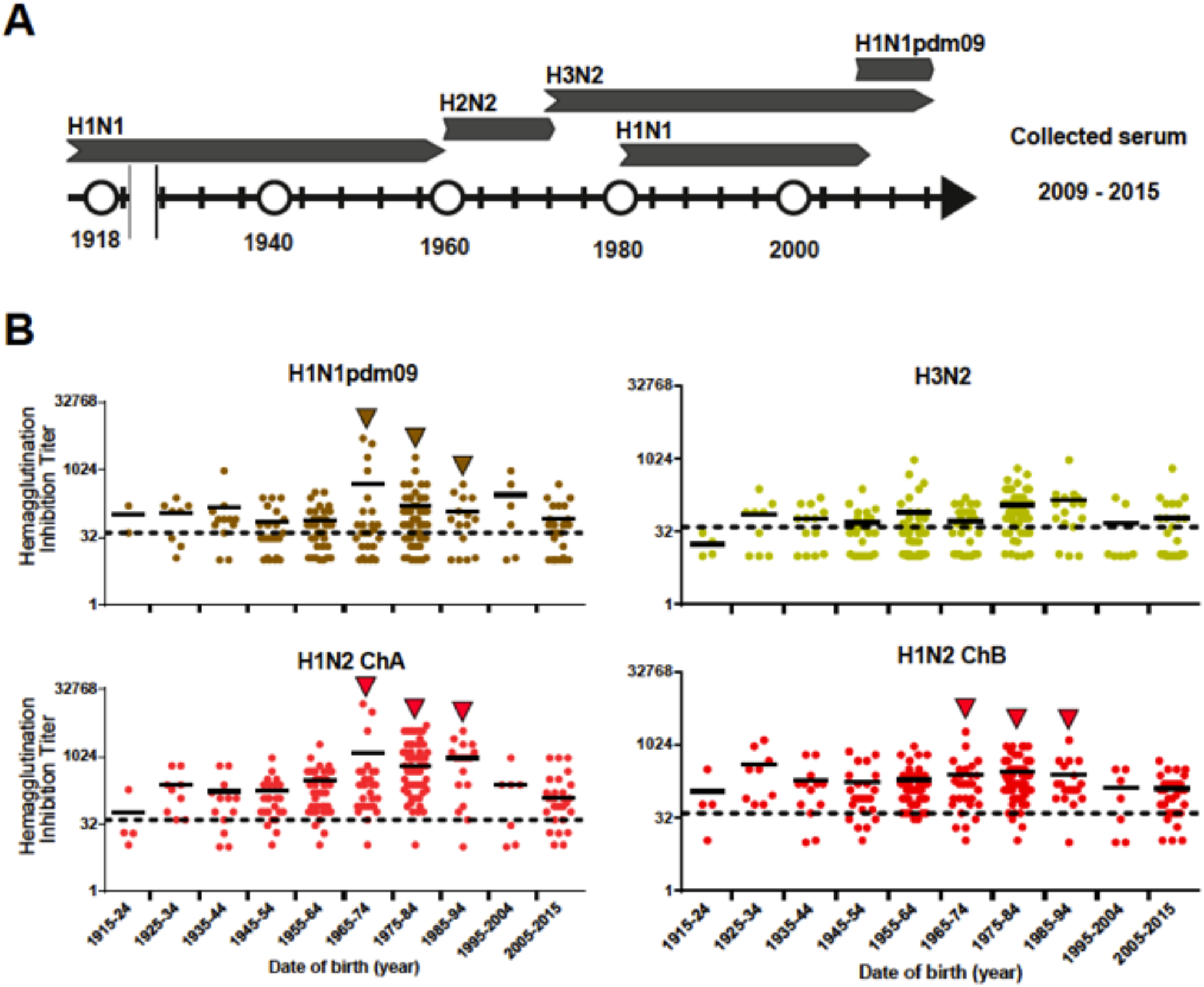
Serological protection of humans of different ages against the Chilean swine IAVs. **(A)** Influenza A strains that circulated in human populations since 1918. H1N1: 1918-1957 and 1977-2009; H2N2: 1957-1968; H3N2: 1968-to date: A(H1N1)pdm09: 2009-to date. **(B)** 237 human sera collected from 2009 to 2015 were evaluated against the Chilean swine viruses by HIA. Serum titers against the A(H1N1)pdm09, H3N2, H1N2 ChA and H1N2 ChB swine viruses are grouped and shown according to individual’s date of birth.

**Figure 11.**
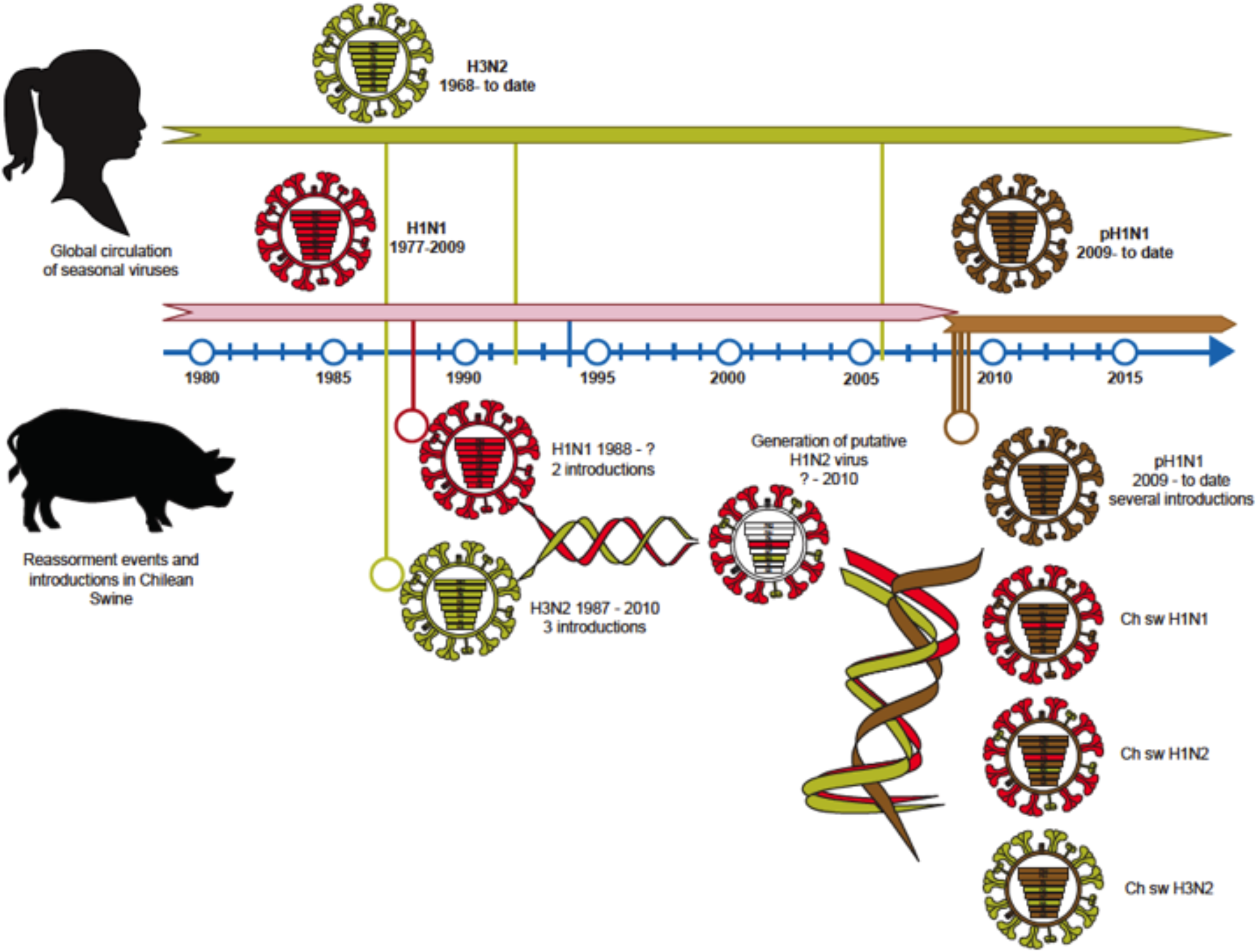
Model of the introduction of human influenza viruses into swine in Chile leading to the generation of novel swine IVAs with zoonotic potential. Human H1N1 and H3N2 seasonal viruses were introduced into swine as early as the mid to late 1980s. The H1N1 were introduced at two time points during this decade, whereas the H3N2 subtype was introduced in the early 1990s and then again sometime after 2005. This likely generated double reassortment events that gave rise to the ChH1N2A and ChH1N2B lineages and to the rare H3N2 viruses found in Chile. Upon the emergence of the 2009 H1N1 human pandemic, this strain was rapidly introduced into the Chilean swine population early during the pandemic and thereafter in multiple occasions generating multiple additional reassortment events that gave rise to the increased diversity of genotypes found in the country. In all cases, all the viruses have the 6 internal genes belonging to the A(H1N1)pdm09 strain, which replaced the previously exiting internal genes (currently of unknown origin).

## Discussion

Here we present the first in depth study of the origin, genetic diversity, and antigenic characteristics of swine IAVs in Chile, an important producer of commercial swine in South America. Using whole genome sequencing and antigenic cartography we show that four IAV lineages are present endemically in swine production farms, ChH1N2A, ChH1N2B, a swine H3N2 and A(H1N1)pdm09-like. Of interest, the H1N2 and the H3N2 viruses were found to originate from human seasonal viruses introduced to the swine population multiple times from as early as the mid to late 1980’s. Our results also show that the introduction of the A(H1N1)pdm09 strain into swine, after the 2009 human pandemic, generated a series of reassortment events that further diversified the swine IAV genotypes (Fig. 2), and completely displaced the internal genes of the pre-pandemic swine IAVs endemic stains (Fig. 11). This has resulted in the genesis of viral clusters that are antigenically distinct, and that differ from other swine IAVs previously reported in Latin America and globally, and hence this suggests that these viruses have evolved independently in Chile. Importantly, given the time of introduction of the older non-pandemic HA and NA viral proteins, our results indicate that the general population (individuals of all ages) is susceptible to the swine H3N2 viruses and that the elderly and young children (<10 years of age) also lack protective antibodies against the swine H1N2 strains, demonstrating that these viruses could be potential zoonotic threats.

Our initial serological analysis of representative production farms revealed a high prevalence of IAV, suggesting its endemic circulation in the swine population (Fig. 1a-d). Analysis of respiratory samples obtained from 8–12-week-old animals confirmed the consistent identification and isolation of swine IAVs in all farms tested. Furthermore, our results indicate that the swine IAVs in Chile have a gene constellation that originated from previous and more recent introductions of human IAVs into swine, resulting in multiple reassortment events over a period of ∼30 years. The introduction of human HA and NA genes took place as early as 1988 for the ChH1N2B viruses and in 1994 for the ChH1N2A strain, and in 1987 and 2006 for the singleton H3N2 viruses, which differ from other swine IAVs described recently in Latin America (Cappuccio et al., 2011; Dibarbora et al., 2013; Gonzalez-Reiche et al., 2017; Mena et al., 2016; Nelson *et al*., 2015b; Nelson *et al*., 2019; Sanchez-Betancourt et al., 2017). Additionally, as reported elsewhere, after 2009 the pandemic human A(H1N1)pdm09 strain was also introduced multiple times generating further reassortment (Howard et al., 2011; Kirisawa et al., 2014; Nelson et al., 2015c; Pereda *et al*., 2010; Tinoco et al., 2016). Overall, three to four putative reassortment events between these viruses generated eight distinct swine IAVs genotypes that circulate in the swine population today (Fig. 2 and 11).

We had previously reported the identification of the H1N2 strains from Chile, however; the in depth characterization of the ChH1N2B and ChH1N2A clusters presented here (also classified as humans seasonal lineage 1B.2 according to the new proposed designation(Anderson *et al*., 2016)) had not been done(Nelson *et al*., 2015a; Tapia et al., 2018; Tapia et al., 2020). The ChH1A virus appears to have spread more efficiently in pigs, since it was found in 12 farms that were geographically unrelated, while ChH1B was only found in two farms (Fig. 1A and 4A). Our results also suggest that the N2 Chilean cluster originates from the introduction of a human H3N2 virus, possibly in the mid-1990s’. This strain appears to have disseminated widely in swine farms, like what was observed for the Chilean H1 clusters (Fig. 6). Importantly, most of the viruses identified (> 50% of sequenced viruses) had an H1 HA and an N2 NA segments corresponding to these older human derived influenza strains, which are not related to IAVs found in swine elsewhere(Nelson *et al*., 2015d; Watson *et al*., 2015). The phylogenies of both gene segments showed long branch lengths (Fig. 4A and 6), which is likely due to lack of swine surveillance data for prolonged periods of time (prior to 2012). However, when compared to human seasonal strains these swine IAVs also clustered separately (Supplementary Fig. 2a and c). This indicates that these viruses have undergone substantial evolution and drift in swine, which was also evidenced when antigenic cartography was performed (Fig. 9).

Other human origin H1 viruses introduced into the swine population in the 1990’
ss have been reported in other Latin American countries (Nelson *et al*., 2015b; Nelson *et al*., 2019). Nevertheless, these viruses are substantially different from the Chilean strains. It must be noted that the lack of IAV surveillance in swine in Chile before 2012, limits our analytic potential to determine the time of circulation and whether the H1 and N2 genes were prevalent among the swine population prior to this year. It is plausible that prior to the introduction of the A(H1N1)pdm09 strain the Chilean swine viruses might have circulated as the original human derived H1N1 and H3N2 subtypes, which eventually generated the H1N2 after reassortment in swine rather than being a direct introduction of human H1N2 viruses that circulated for very limited period, as was suggested in the case of the Brazilian swine H1N2 strains. Additionally, the ChN2 sequences were more closely related to N2 segments present in human H3N2 viruses from the 1990’s. Hence, this further supports the notion that the ChH1N2 viruses are swine reassortant viruses. Nevertheless, additional swine IAV sequences from samples obtained prior to the 2009 pandemic would be necessary to evaluate this further.

Our study also revealed that the A(H1N1)pdm09-like viruses circulate endemically and as a predominant strain in commercial swine. The topology of the tree suggests that there has been seven independent human-to-swine introductions of the A(H1N1)pdm09 strain since 2009. Noteworthy, two swine viruses clustered closely together with human seasonal A(H1N1)pdm09 viruses from Chile sequenced a year earlier (Fig. 4a and 5b), demonstrating that recent and constant reverse zoonotic events contribute to the dynamics and genetic diversity of swine IAVs in Chile. Phylogenies also demonstrated that internal segments from all the 52 whole genome sequences belonged to the A(H1N1)pdm09-like lineage (Fig. 7 and 8, and Supplementary Fig. 1a-d). Reassortment of internal A(H1N1)pdm09-like genes resulting in IAVs with either a mixture of pandemic or endemic glycoproteins, have also been observed and reported globally (Mena *et al*., 2016; Nelson *et al*., 2015c; Vijaykrishna et al., 2010; Vijaykrishna et al., 2011; Zhu et al., 2011), including a predominant reassortant Eurasian avian-like (EA) H1N1 virus, that contains A(H1N1)pdm09-like and triple-reassortant (TR)-derived internal genes that has been found in swine since 2016 in China(Sun et al., 2020). Nonetheless, the complete displacement of previous internal genes by the A(H1N1)pdm09-like genes in all viruses detected in a surveillance study is highly unusual and has not been reported in other regions (e.g. in Eurasian, Asia and the United States), where classical or avian-like lineages that existed prior to 2009 are still prevalent(Howard *et al*., 2011; Nelson *et al*., 2015c; Watson *et al*., 2015; Yang et al., 2016; Zhu *et al*., 2011).

Our phylogenetic analyses showed that among the pandemic internal genes, the M segment, except for three sequences, formed a monophyletic cluster regardless of whether it belonged to the A(H1N1)pdm09-like, swine H3N2 or the ChH1N2A and B strains (Fig. 7). Previous studies have suggested that the M segment of the pandemic strain contributed to its increased transmissibility (Chou et al., 2011; Lakdawala et al., 2011). Hence the non-random reassortment and incorporation of this specific M gene, derived from an early 2009 A(H1N1)pdm09 human virus, suggest that this gene is well-adapted in swine and probably provides a replication advantage in this host allowing this M segment to almost outcompete any other M genes completely. This is highly relevant from a public heath perspective since H3N2 virus variants containing the A(H1N1)pdm09 M segment have been responsible for zoonotic transmission from swine to human in the US (Epperson et al., 2013; Greenbaum et al., 2015). Hence, a further phenotypic analysis of this M segment is warranted to understand its potential contribution to the emergence of zoonotic viruses.

In contrast, the other internal genes showed evidence of multiple introductions suggesting a high level of reassortments and increased divergence dynamic since the introduction of the A(H1N1)pdm09 strain (Fig. 8, and Supplementary Fig. 1a-d). Of note, the NS segments demonstrated a unique origin for each farm. Interestingly, there were no genotypes in common among farms when all genes are considered. This suggests that there is poor or no recent transmission among farms. However, due to the limited number of IAV isolates per farm, we cannot confirm this in the current study and further long– term epidemiological studies would be needed.

Of high interest, we identified two different H3N2 viruses corresponding to different singletons in the H3 and N2 phylogenies (Fig. 3b). A similar H3 gene has been previously described and was found to be related to other South American viruses(Nelson *et al*., 2015a). We estimated that this strain was introduced in ∼2006 (Fig. 3b). However, the other H3 virus identified was distantly related to the H3 cluster I and had a TMRCA estimated at around 1987. In the case of the N2 segments, we found evidence of four different introductions of these genes into swine. One major cluster, representing 89% of the N2 genes sequenced, corresponds to viruses introduced in the early 1990’s, which appeared to have evolved in swine since then but did not spread to other farms (Fig. 6). However, we found three additional N2 singletons. One of them corresponds to an older introduction of a human H3N2 in the 1980’s, while the other two N2 genes correspond to contemporaneous human viruses, which were likely introduced sometime in the late 1990’s to early 2000’s. The identification and isolation of these minor circulating H3N2 strains in swine, even after >30 years since their introduction is remarkable and suggest that swine IAVs can evolve and be maintained for prolonged periods of time. Surprisingly, these viruses were found to be substantially different to all human and swine H3N2 viruses (Fig. 3b, 6 and Supplementary Fig. 2b), indicating that they are unique and could be potential zoonotic threat to naïve humans.

Our antigenic characterization revealed that the ChH1N2A and B strains are antigenically divergent from each other (Fig. 9). Interestingly, these analyses also showed the presence of antigenic drift variants of the Chilean swine A(H1N1)pdm09-like and the ChH1N2 strains. Given that most farms use poorly antigenically matched (non-homologous) influenza vaccines or do not vaccinate at all, it becomes highly relevant to further understand the evolutionary dynamics of swine IAVs in Chile, and to determine the factors that are driving the antigenic drift in these farms. The vast genomic and antigenic differences observed in these viruses and their divergence from existing vaccines (Fig. 9 and Supplementary Fig. 2a-c), prompted us to conduct a risk assessment to determine their zoonotic potential(Cox et al., 2014). Serological analyses from individuals of different ages indicated that an important proportion of the population is likely to be susceptible to the swine H3N2 strain, and that there is a likelihood of moderate protection against the ChH1N2A, ChH1N2B and swine A(H1N1)pdm09-like viruses. Importantly, most of the individuals that show protective titers were born during the period of circulation of the human strains from which these swine influenza viruses are derived (Fig. 11). This supports the concept that the first encounter with the virus has a strong effect on the antibody imprinting of individuals (Nachbagauer et al., 2017), and highlights the notion that with the passing of time new naïve individuals become susceptible to IAVs maintained in swine. Additionally, we identified the swine H3N2 strain as higher zoonotic risk potential to humans, and hence this emphasizes the need for continuous long-term surveillance of swine IVAs in the region.

To date this is the largest molecular epidemiological study of swine IAV in South America. Our results demonstrate that IAV is widespread in commercial swine in Chile, revealing three novels antigenically distinct IAVs, a ChH1N2A, a ChH1N2B and swine H3N2 strains, with the later virus posing a potential zoonotic risk due to the poor immunity found in the general population. Overall, this study contributes to the understanding of the global and regional circulation of swine IAVs, the dynamics of swine IAV transmission in production farms and to the design of long-term surveillance and prevention strategies.

## Materials and Methods

### Swine study design and sample collection

All animal procedures were approved by the Institutional Animal Care and Use Committees of both, Universidad de Chile (protocol numbers 20-2015 and 02-2016) and Pontificia Universidad Católica de Chile (PUC, protocol number 13-001).

Commercial swine farms were invited to participate in an active surveillance program for the identification swine IAV. Enrolled herds were representative of modern swine production systems and included breeding herds, nurseries, finishers and wean-to finish farms. To determine the age at which swine became infected with swine IAV, in 2013 we performed a serological study in three farms. On one farm, we conducted a longitudinal study where 47 animals were tested periodically to determine the seroconversion rate at different time points of during the different production stages (farm 1 on Fig. 1). On two additional farms we conducted a cross-sectional study in which animals from all ages were sampled in a single day (farms 2 and 3, Fig. 1). Serum samples were directly provided by the local veterinarian at each production site. During December 2013 to January 2015, samples were obtained during a total of 55 visits to 39 farms (24 companies) comprising 93.8% of the commercial farms in Chile. Farms were in the central regions of Chile (Metropolitan, Valparaíso, O’Higgins, Maule, Biobío, and Araucanía Regions), where ∼99% of the intensive pig production is found (Neira et al., 2017b). Due to confidentiality agreements the geographical locations of the farms are not disclosed. During each visit, nasal swabs and oral fluids were collected from pigs 6 to 14 weeks of age which is the age when there is an increased probability of influenza detection (Corzo *et al*., 2013). Samples were obtained from pigs with influenza-like clinical signs, which included coughing, sneezing, fever and/or lethargy. If no animals were found with influenza-like signs during the visit, samples were obtained randomly. Nasal swabs collections consisted in selecting 30 animals, which were manually restricted, and samples were obtained from both nostrils using nylon flocked swabs that were placed in tubes with 1 ml of universal viral transport media(Esposito et al., 2010). The sample size per visit was estimated to detect at least one IAV positive sample assuming ≥10% prevalence of IAV in the farm and considering 95% confidence of viral detection. Additionally, approximately 6 oral fluid samples were obtained from swine of the same age using 1 m of twisted cotton rope hanging for 20 min in each pen. Ropes were squeezed into a sterile plastic bag and the oral fluids were deposited into a sample tube(Romagosa et al., 2012). Additional samples were also submitted for diagnostics (including lung tissue) when suspected cases of IAV were observed. Samples collected were refrigerated and immediately transported on ice to the laboratory for processing.

### Diagnostic tests

To determine generic anti-IAV antibodies, 50 μl serum samples were tested in using a commercially available multistep competitive influenza virus-specific ELISA kit that tests for antibodies against the Nucleoprotein (IDDEX, Westbrook, Maine, USA). Oral fluids were centrifuged at 5,000 rpm for 20 min and the supernatant was stored in aliquots at - 80°C until further processing. Both nasal swabs and oral fluids were tested for IAV using real time RT-PCR (rtRT-PCR) and/or virus isolation (VI). For rtRT-PCR, viral RNA was first extracted using the TRIzol® LS Reagent (Invitrogen™, Carlsbad, CA, USA) according to the manufacturer’s instructions, and then followed by amplification of the matrix gene using a previously described protocol for IAV diagnosis (Organization, 2011). rtRT-PCR results with cycle threshold (Ct) < 35 were considered positive.

Viral isolation was attempted in rtRT-PCR positive samples. This test was carried out in Madin-Darby Canine Kidney (MDCK) cells using minimum essential medium (MEM) supplemented with 10% fetal calf serum (FCS) and 1% antibiotic-antimycotic solution (Organization, 2011). Briefly, the MDCK monolayers were washed twice to remove FCS using PBS containing 1 μg/mL of trypsin treated with N-tosyl-L-phenylalanyl chloromethyl ketone (TPCK) (Sigma-Aldrich, St. Louis, Mo, USA), inoculated with each sample, and incubated for virus absorption for 1h at 37ºC. Subsequently, cells were rinsed with PBS to eliminate unbound virus, and IAV growth medium (MEM supplemented with 1 μg/mL of TPCK trypsin, 0.3% bovine serum albumin, and 1% antibiotic-antimycotic solution) was added. The monolayers were incubated at 37°C. Each sample was observed for cytopathic effect (CPE) daily for up to 5 days. Plates without CPE were passaged again and observed for another 5 days. Samples with no CPE after the second passage were considered negative for IAV. CPE positive samples were tested by hemagglutination assay using turkey erythrocytes and rtRT-PCR to confirm the presence of IAV.

### Viral sequencing of swine samples

IAV isolates were sequenced by whole genome sequencing and in a subset of samples (18 isolates) the HA gene was sequenced by Sanger. For whole genome sequencing, the samples were prepared using a multisegment RT-PCR (mRT-PCR), following the protocol described by Zhou et al. (2009)(Zhou et al., 2009). Eluted RNA from each positive sample was used as template to synthesize cDNA using primer MBTuni-12 (5’-ACG CGT GAT CAG CRA AAG CAG G-3’) and Superscript III First Strand Synthesis SuperMix (Invitrogen, Carlsbad, CA, USA), following the manufacturer instructions. Then the cDNA was amplified by 5 cycles (94°C for 30 s, 45°C for 30 s, and 68°C for 3 min), and then 31 cycles (94°C for 30 s, 57°C for 30 s, and 68°C for 3 min). The total volume for the RT-PCR reaction was 50 μl using a High-Fidelity DNA Polymerase (Agilent, Santa Clara, USA), with MBtuni-12 and MBtuni-13 (5’-ACG CGT GAT CAG TAG AAA CAA GG-3’) primers (buffer: 5 μl, 10mM dNTPs: 1 μl, 1.5 μl of each primer, Picomax Enzyme: 1μl, water: 35 μl, and 5μl of cDNA). PCR products were run in a 1.5% agarose gel by electrophoresis, and purified using QIAquick Spin Kit (QIAGEN, Valencia, CA, USA). Purified PCR products with ≥ 25 ng/μl of DNA concentration were submitted to the Center for Research on Influenza Pathogenesis (CRIP) Sequencing Core at the Icahn School of Medicine at Mount Sinai, for library preparation and sequencing using next generation sequencing (NGS) technologies (Illumina HiSeq2000). Illumina reads were mapped to an IAV reference genome using Bowtie2 and consensus sequences extracted using SAMtool. The HA Sanger sequences were obtained from IAV isolates submitted for diagnosis to the Veterinary Diagnostic Laboratory at University of Minnesota (VDL-UMN) during 2012. All the whole genome and HA sequences were deposited in GenBank (Supplementary Table 1) and were included in the phylogenetic analyses.

### Human sample collection and A(H1N1)pdm09 virus sequencing

All the patient and volunteer samples used in this study were collected after written informed consent was obtained under protocols 11-116 and 09-203, which were reviewed and approved by the Scientific Ethics Committee of the School of Medicine at Pontificia Universidad Católica de Chile (PUC) before the start of sample collection. Viral isolates prior to 2009-2011, were obtained anonymously through the Clinical Diagnostic Laboratory of the UC-Christus Health Network as part of their seasonal epidemiological surveillance.

To elucidate the diversity and time of introduction of the A(H1N1)pdm09-like viruses, we sequenced the full genome of 114 A(H1N1)pdm09 viruses obtained from infected individuals during 2009 to 2013 in Chile. Viral RNA was isolated using a ZR 96 Viral RNA kit (Zymo Research). The IAV genomic RNA segments were simultaneously amplified from 3 μl of purified RNA using the mRT-PCR described above (Zhou *et al*., 2009; Zhou and Wentworth, 2012). The influenza genome amplicons were barcoded and amplified using an optimized sequence-independent single primer amplification (SISPA) protocol(Djikeng et al., 2008; Djikeng et al., 2009). Subsequently, the SISPA amplicons were purified, pooled and size selected (∼800 or∼200 bp), and the pools were used for library construction and the complete coding genomes of the IAVs were then sequenced using a high-throughput next-generation sequencing pipeline at the J. Craig Venter Institute using the 454/Roche GS-FLX platform using Titanium chemistry or the Illumina HiSeq 2000 platform. The sequence reads were sorted by barcode, trimmed, and searched by TBLASTX against custom nucleotide databases of full-length IAV segments downloaded from GenBank to filter out both chimeric influenza sequences and non-influenza sequences amplified during the random hexamer-primed amplification. The reads were binned by segment and the 454/Roche GS-FLX reads were *de novo* assembled using the clc_novo_assemble program (CLC Bio). The resulting contigs were searched against the corresponding custom full-length Influenza segment nucleotide database to find the closest reference sequence for each segment. Both, the 454/Roche GS-FLX and Illumina HiSeq 2000 reads were then mapped to the selected reference IAV segments using the clc_ref_assemble_long program (CLC Bio). At loci where both 454/Roche GS-FLX and Illumina HiSeq 2000 sequence data agreed on a variation (as compared with the reference sequence), the reference sequence was updated to reflect the difference. A final mapping of all next-generation sequences to the updated reference sequences was then performed. Any regions of the viral genomes that were poorly covered or ambiguous after next-generation sequencing were amplified and sequenced using the standard Sanger sequencing approach.

### Phylogenetic analysis

Reference human and swine sequences from different known influenza lineages, and the most similar sequences available in GenBank obtained using the BLAST tool from NCBI (http://blast.ncbi.nlm.nih.gov/Blast.cgi) were used to reconstruct the genetic phylogeny of the Chilean swine viruses. Sequences of each of the segments were aligned using MUSCLE and then a Bayesian evolutionary analysis was performed by sampling three methods using the BEAST 1.8.2 software (Drummond et al., 2012). We used the Partition finder software to select the substitution model (Lanfear et al., 2012) and codon partition with the lowest Bayesian information criteria, and used the uncorrelated exponential clock model and a coalescent Bayesian skyline tree, prior to running the analyses. Each segment phylogeny was run for 500 million generations, sampling every 50,000 trees. Convergence and mixing of the simulations were assessed using Tracer, and we then used Figtree v1.4.2 (Vrancken et al., 2015) to visualize the maximum clade credibility tree (MCC). TMRCA and substitution rates for specific branches were obtained from the MCC. Trees were illustrated highlighting Chilean swine, Chilean human, South American swine, and other reference sequences from human and swine as shown. For the phylogenetic analyses we used an alignment of 296 sequences of the H1pdm09 viruses, 156 H1 sequences with viruses from the delta swine cluster and human seasonal strains, and 261 H3 sequences including the viruses sequenced in this study. Alignments of sequences obtained in this study and reference sequences used to reconstruct the phylogeny of internal genes comprised: 189 sequences for PB2, 188 sequences for PB1, 198 sequences for PA, 163 sequences for NS, 204 sequences for NP, 210 sequences for NA (N2), 259 sequences for NA (N1pdm09), and 205 sequences for M.

To determine the phylogenetic relations of the novel Chilean swine viruses, seasonal human viruses and human vaccine stains were used with a total of 185 H1 and 173 H3 HA sequences from GISAID/GenBank databases to make backbone trees for each H1 and H3 swine lineages. In addition to the backbone trees, 41 H1 and 2 H3 swine Chilean sequences were added to the tree for phylogenetic analysis. Data were aligned via MUSCLE (Edgar, 2004) and sequences were trimmed to the beginning of mature H1 HA protein gene sequence using BioEdit v7.0 (Hall, 1999). Approximate maximum likelihood trees (Juke-Cantor model) were constructed using the Mega 7.0 software package (Tamura et al., 2011). Gene sequences obtained from this study were submitted to GenBank and GISAID under accession numbers in Supplementary Table 1.

### Antigenic analysis of H1 swine IAVs

To elucidate whether the level of mutations observed also resulted in changes in the antigenic properties of the viruses, we evaluated the antigenic cross-reactivity between the swine H1ChA, H1ChB and H1pdm viruses circulating in Chile using antigenic cartography (Smith et al., 2004). First, we selected representative swine Chilean IAVs strains within each phylogenetic cluster and obtained the consensus amino acid sequence of the antigenic sites on the HA gene (Sa, Sb, Ca1, Ca2 and Cb) using Jalview v2.10.0. Distance matrices were then performed between the amino acid sequences of each cluster and their respective consensus sequence using MEGA v6.0. The strains with the highest amino acid identity to the consensus sequence, and those representing different farms and geographical regions, were selected to produce monovalent vaccines. Each selected virus, containing at least 128 hemagglutination units (HAU)/50 μl, was inactivated with 0.1% formalin.

Antisera were produced using pathogen-free four-week-old female guinea pigs obtained from the Chilean Institute of Public Health, which were housed at the animal facilities of the College of Veterinary Medicine at the Universidad de Chile. 200 μl of each vaccine complemented with complete Freund’s adjuvant was administered subcutaneously to four animals, which was followed by a second dose complemented with incomplete Freund’s adjuvant 14 days later. After deep anesthesia, by the intramuscular administration of a mixture of ketamine (30 mg/kg) and xylazine (2 mg/kg), guinea pigs were bled via cardiac puncture on day 28 post first vaccination, at which time animals were euthanized with thiopental sodium (120 mg/kg) administered intraperitoneally. Seroconversion was evaluated by hemagglutinin inhibition assay (HIA), following a standard protocol (Kitikoon et al., 2014). Positive antisera with a homologous HIA titer of ≥640 were used for subsequent analyses. Briefly, viruses were titrated to hemagglutination titer of 80 HAU/50 μl and were then tested against homologous and all available heterologous viruses. An HIA titer ≥ 20 was considered positive. The HIA results were used to determine antigenic clusters using the antigenic cartography analysis tool (http://www.antigenic-cartography.org/) (Smith *et al*., 2004).

### Analysis of human antibody cross-protection against H1 swine IAVs

HIA was performed as above to determine the cross-reactive antibody titers in the general population against the ChH1N2s, H1N1 and H3N2 swine IAVs. Between July 2009 and December 2015, a total of 237 blood samples were collected from individuals diagnosed with IAV in Santiago, Chile. Serum samples were obtained by centrifugation at 3000 rpm for 5 min and were then inactivated by trypsin-heat-periodate treatment. Antibody titers against the Chilean swine viral strains were determined by the HIA as previously described (Manicassamy et al., 2010).

## Acknowledgments

We thank the staff of the Instituto de Salud Pública de Chile for biological supplies. We are grateful to Daniela Jimenez, Victoria Garcia, Miguel Vivar and swine veterinarians for all their support in technical assistance and sample collection. We also thank Charles Davis for his support in the phylogenetic risk assessment against human vaccines.

The findings and conclusions in this report are those of the authors and do not necessarily represent the views of the Centers for Disease Control and Prevention or the Agency for Toxic Substances and Disease Registry.

## Funding

Comisión Nacional de Investigación Científica y Tecnológica de Chile (CONICYT) FONDECYT 1161791 (RAM)

CONICYT, FONDECYT 11170877 (VN)

CONICYT, FONDECYT 1211517 (VN)

Fondo de Investigación Veterinaria, FAVET UCHILE 2017 (V.N.)

CONICYT, Programa Beca Doctorado Nacional (RT, JM)

CONICYT, Anillo PIA ACT 1408 (RAM, VN)

Center for Research in Influenza Pathogenesis (CRIP), a National Institute of Allergy and Infectious Diseases–funded Center of Excellence in Influenza Research and Surveillance (CEIRS), contract number HHSN272201400008C (RAM, VN)

## Author contributions

Conceptual: RAM, VN

Funding: RAM, VN

Sample collection: VN, RAM, MT

Swine surveillance analyses: RT, VN, VG, JM, MS, KT, RAM

Antigenic diversity: RT, VN, VG, JM

Human surveillance analyses: RAM, KT, TG, RR, GB, DW, YS

Phylogenetic analysis: BB, YS

Data analysis: VN, RAM, RT, MS, TG, BB

Writing—original draft: RAM, VN, RT, BB

Writing—review & editing: RAM, VN, RT, BB

## Data and materials availability

All data are available in the main text or the supplementary materials.

## Supplementary Materials

### Supplementary Materials for

**Fig. S1.**
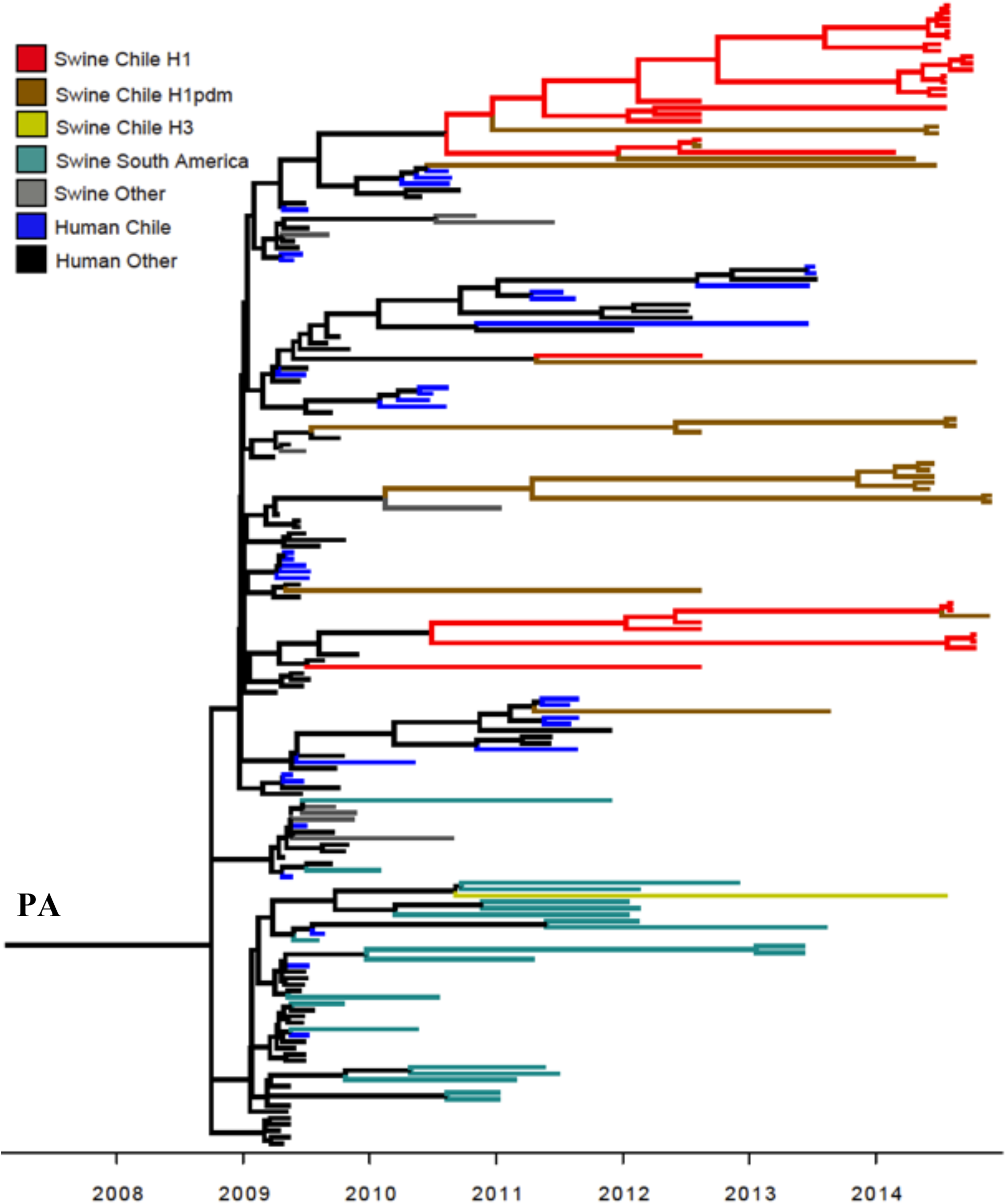
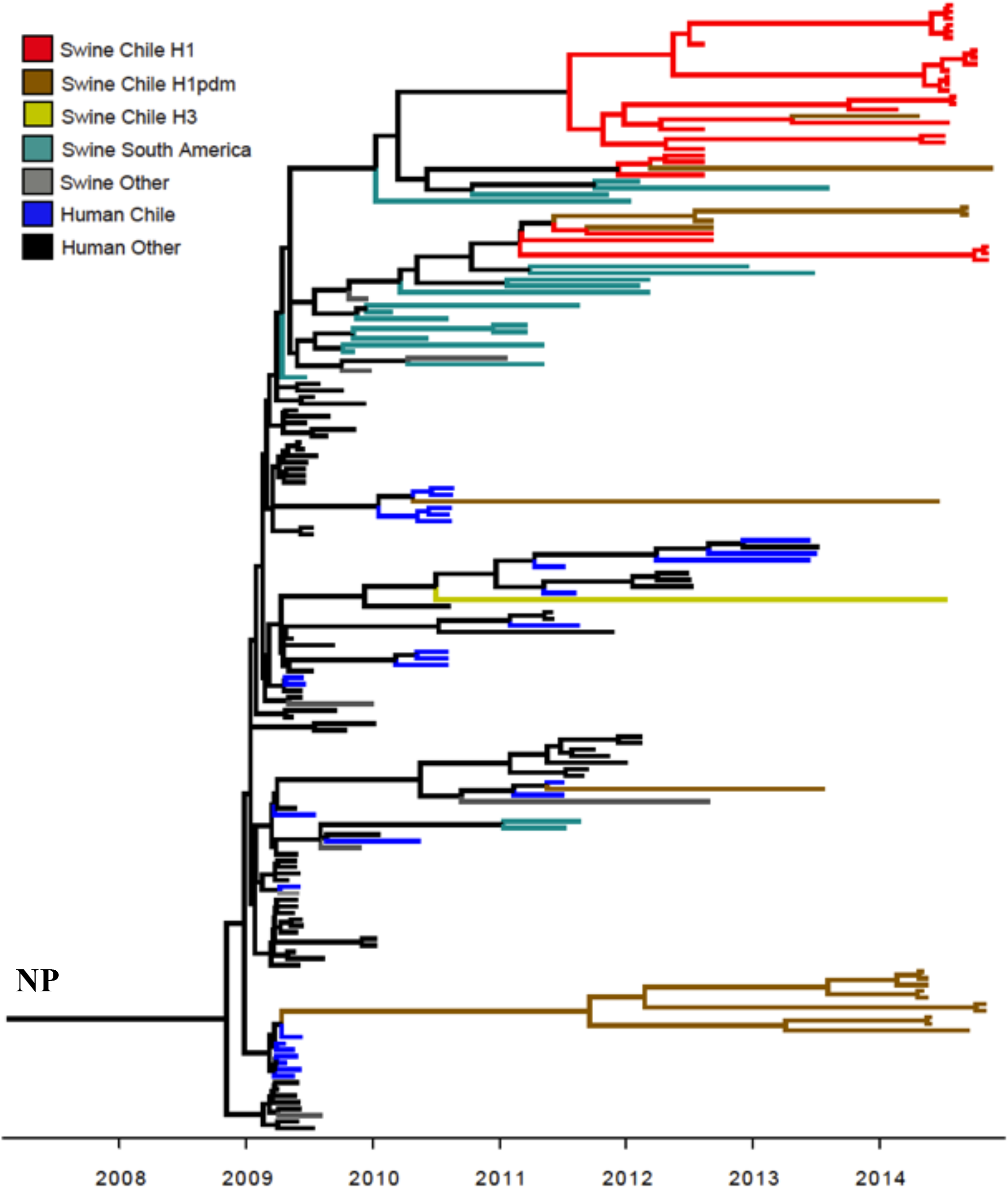
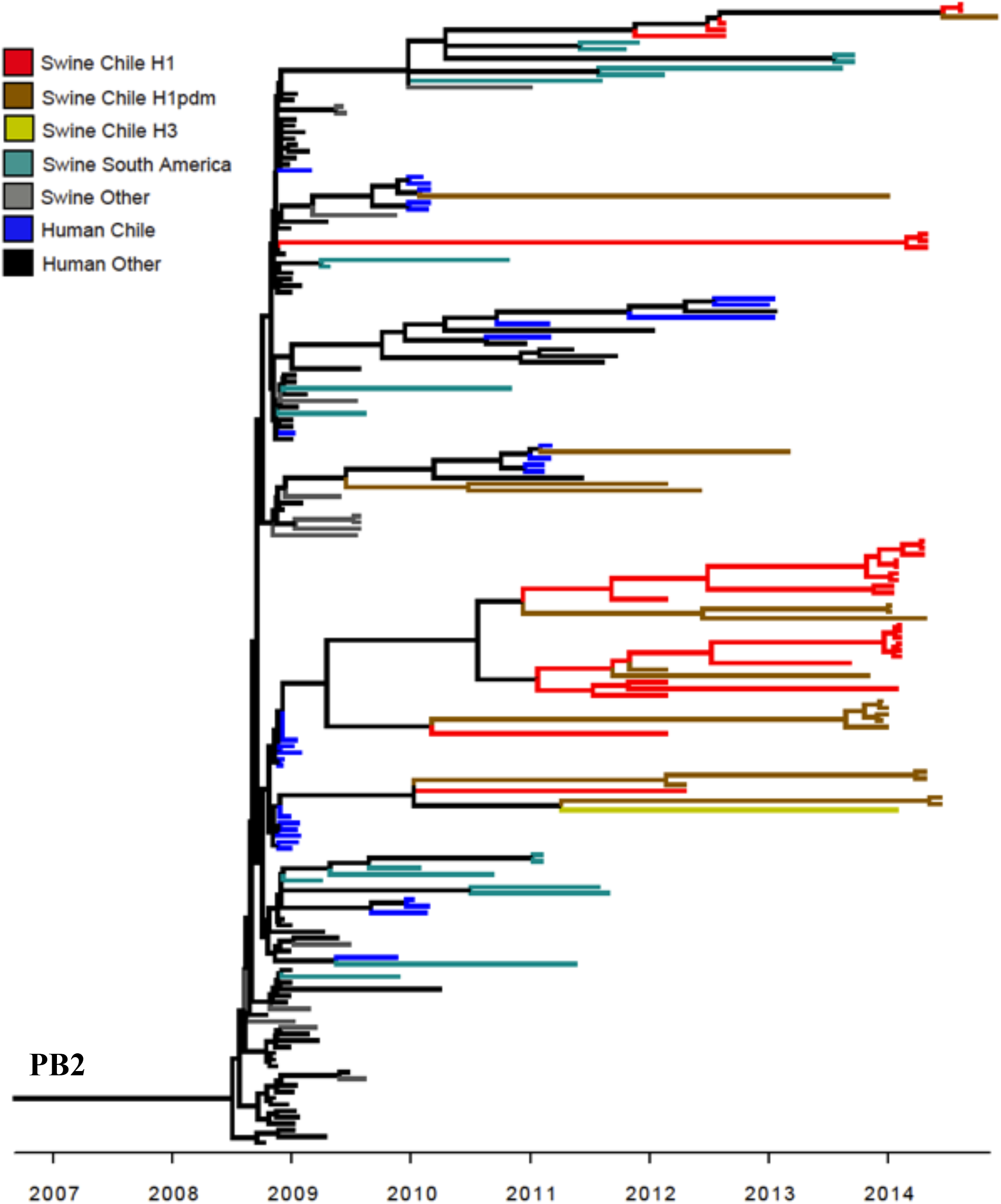
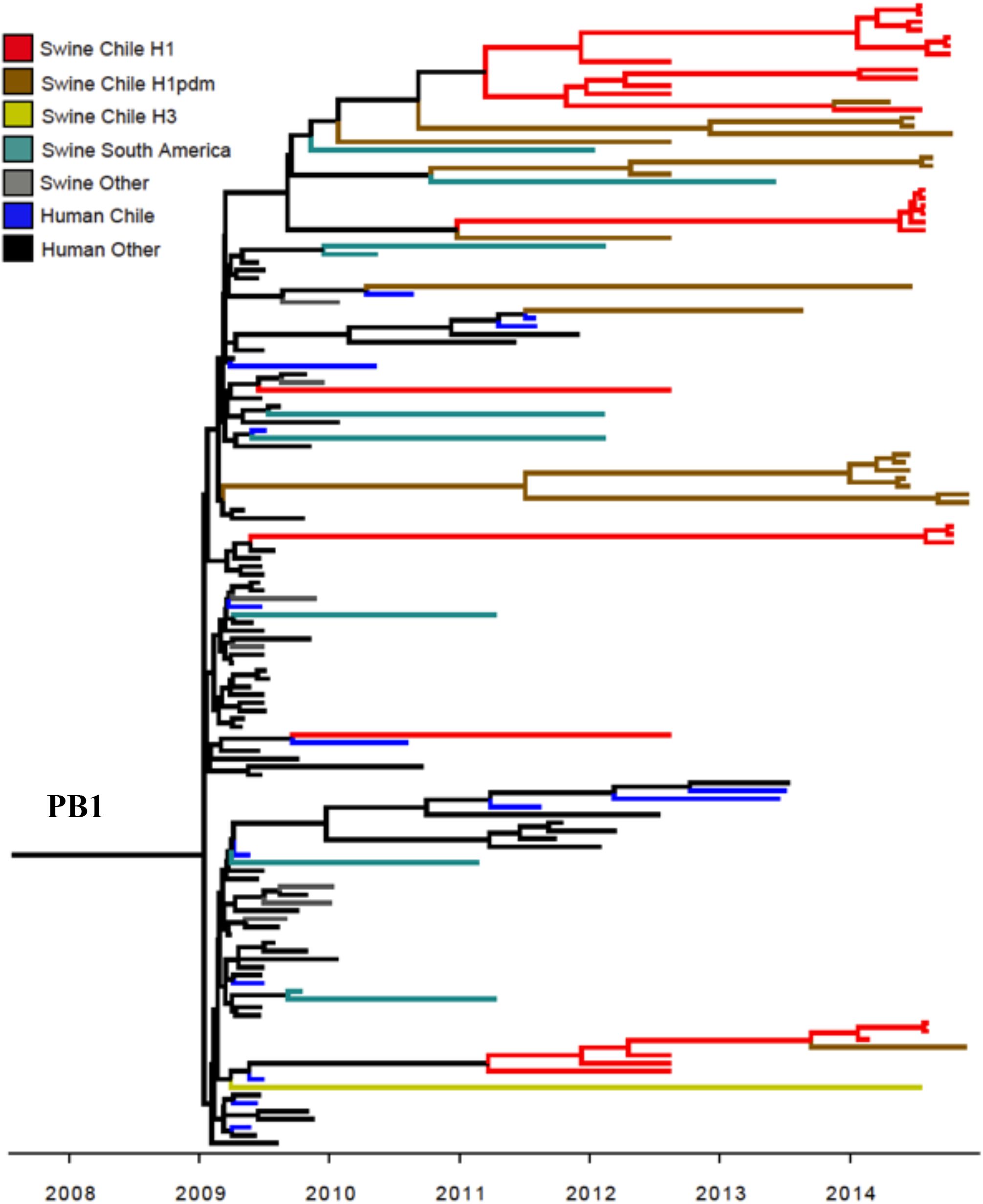
Multiple introductions of the A(H1N1)pdm09 ribonucleoprotein complex genes into the Chilean swIAVs. Phylogenetic relationships of the **(A)** PA, (B) NP, (C) PB2, and (D) PB1 gene segments. Maximum clade credibility tree reconstructed with influenza viruses collected from human and swine. Tree branches are colored according to the HA lineages identified in Fig. 2-4. Bottom axis denotes years as dated in the analyses.

**Fig. S2.**
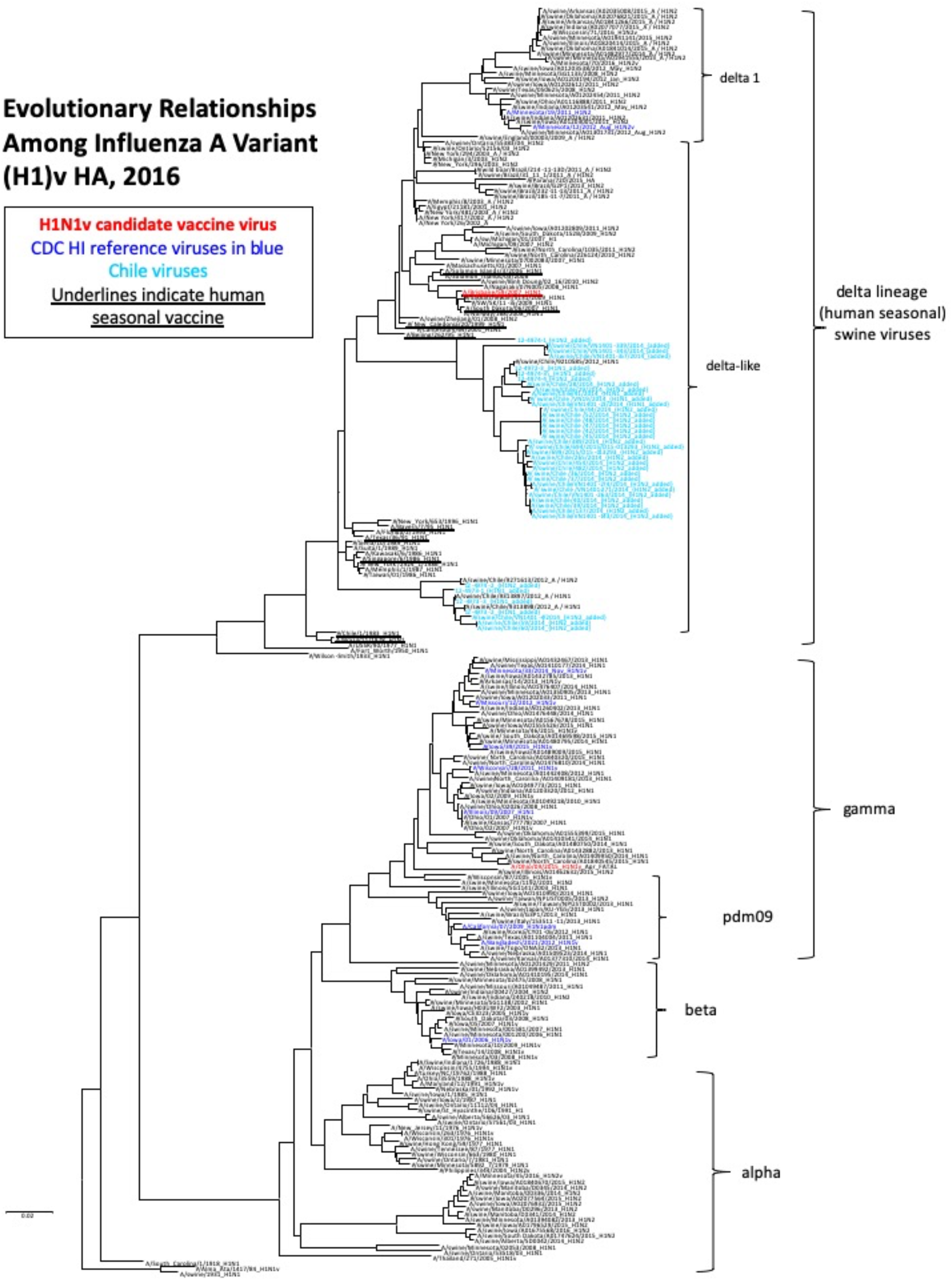
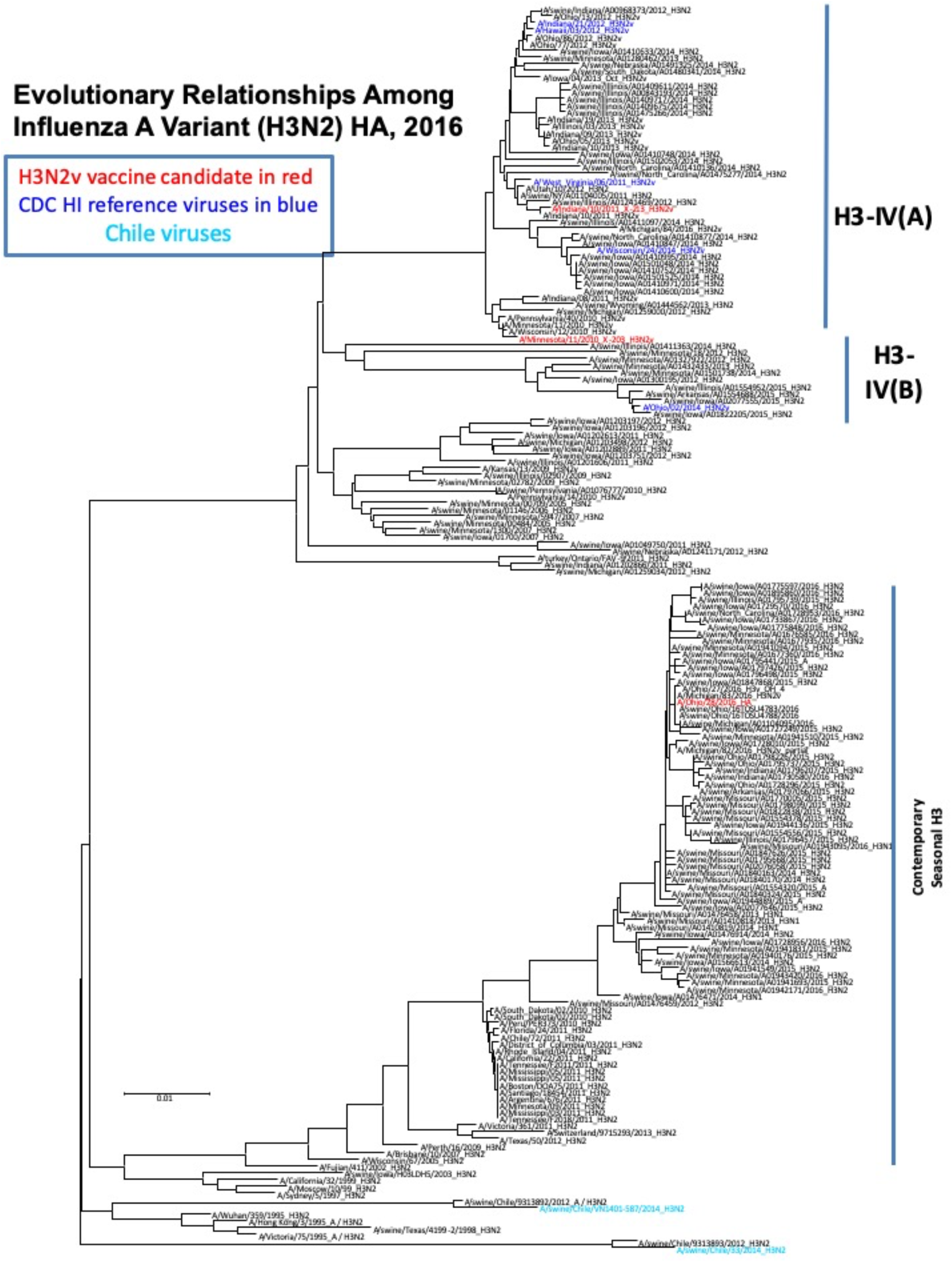
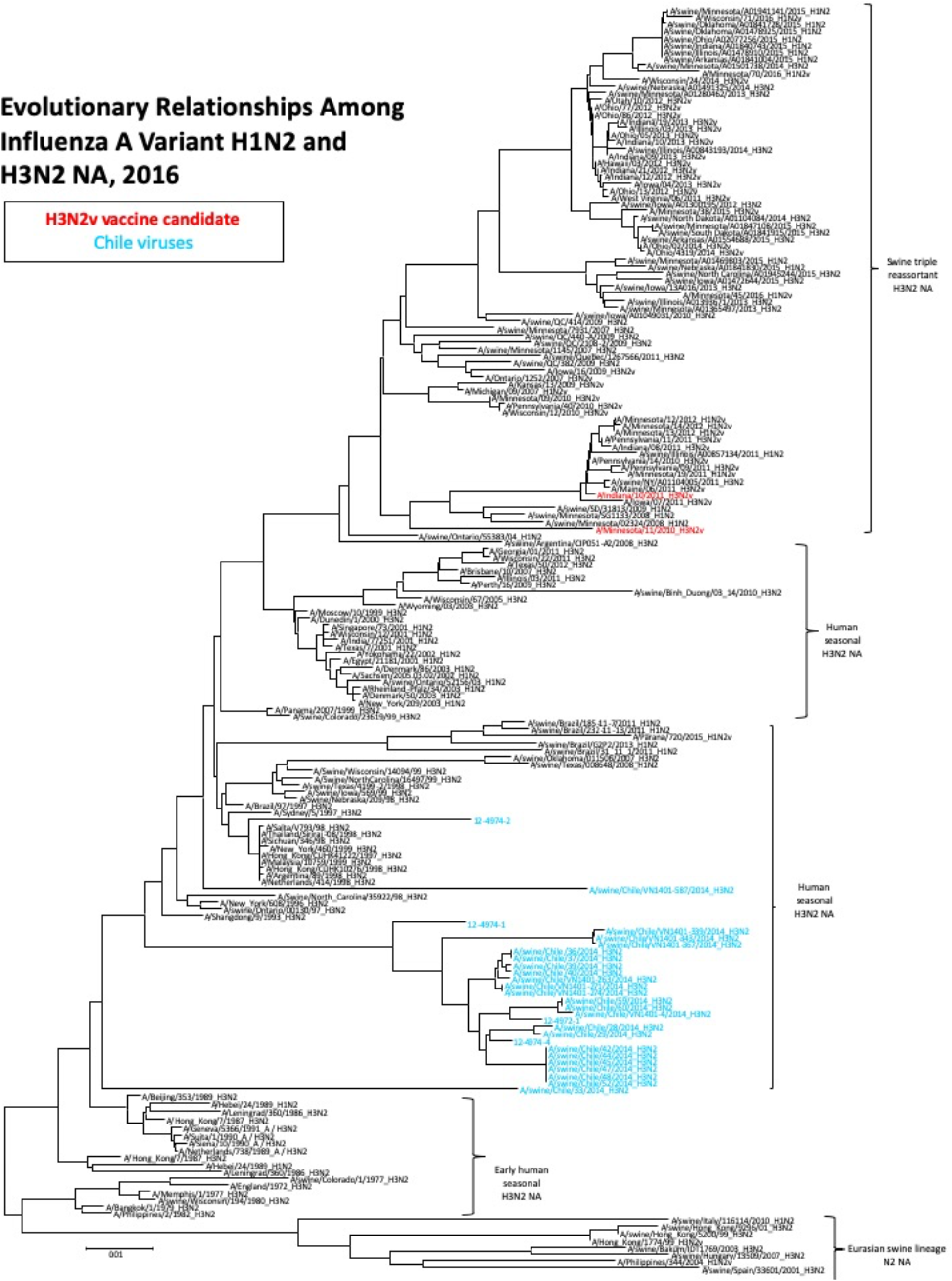
Evolutionary distant relationships among Chilean swIAV variants and vaccine strains. Maximum likelihood tree of the Hemagglutinin genes of H1 (**a**), H3 (**b**) and the N2 gene of H1N2 and H3N2 (**c**) swine lineages. Swine Chilean viruses are in cyan, variant candidate vaccine viruses are in red, and reference viruses of hemagglutinin inhibition by CDC are in blue. Human seasonal vaccine strains are underlined. The scale bar represents nucleotide substitutions per site.

